# *TGFB1*-Engineered Induced Mesenchymal Stromal Cells Exhibit High Immunomodulatory Potency and Durably Reprogram Synovial Macrophages in Osteoarthritis

**DOI:** 10.64898/2026.09.24.753893

**Authors:** Kevin Fan, Mozhgan Rasti, Rebekah Khan, Atoosa Ziyaeyan, Aida Feiz Barazandeh, Rachel Low, Johana Garcia, Shahrzad Nouri, Setareh Safara, Kevin P. Robb, Rajiv Gandhi, Sowmya Viswanathan

## Abstract

Clinical translation of mesenchymal stromal cell (MSC) therapies derived from primary tissue sources is limited by manufacturing inconsistencies driven by donor-to-donor variability, which reduces predictability of clinical response. Induced pluripotent stem cell (iPSC)-derived MSCs (iMSCs) offer a promising donor-independent, clonal alternative strategy that is also more amenable to engineering, but whether engineered iMSCs reproduce the potency attributes of primary MSCs remains unclear. Here, we benchmarked a doxycycline-inducible, hTERT-immortalized iMSC line engineered to overexpress *TGFB1* (*TGFB1-*iMSCs), a key regulator of macrophage polarization, against multiple adipose tissue-derived MSC (MSC(AT)) donors across predefined immunomodulatory and angiogenic potency attributes. *TGFB1*-iMSCs were smaller and more circular with comparable or higher proliferative rates than primary MSC(AT) donors. Interestingly, they had a distinct angiogenic signature (*EDIL3, EDN1, PDGFA*), and nine differentially expressed microRNAs were predicted to target secretory trafficking and growth factor receptor signalling. Functionally, however, *TGFB1-*iMSCs secreted less VEGF and showed intermediate HUVEC tube formation, but matched or exceeded all primary MSC(AT) donors in a monocyte-macrophage transwell immunomodulatory assay. Notably, in a murine destabilization of the medial meniscus (DMM) model of post-traumatic osteoarthritis, a single intra-articular injection of *TGFB1*-iMSCs, but not MSC(AT), reduced total synovial macrophage numbers and lowered the MHCII/CD206 ratio at eight weeks, although it did not reduce cartilage degeneration, synovitis, or fibrosis. Together, these findings show that *TGFB1*-engineered iMSCs have a macrophage-directed potency profile distinct from that of primary MSC(AT). Immortalized iMSCs therefore provide a reproducible platform for mechanistically dissecting engineered MSC potency and its effects on the immune environment.

## Introduction

Despite decades of research, there are only a handful of approved mesenchymal stromal cell (MSC) products (1), including only one approved by the Food and Drug Administration (FDA), after multiple decades of product improvement and clinical trials. Several reasons have been speculated for this failure (2), including heterogeneity in recipient disease stage and baseline inflammatory and immune state (3), which necessitate disease-specific patient stratification (1). Consistency of the MSC product itself has also been identified as a key variable (1, 4, 5, 6), representing an amalgamation of multiple layers of heterogeneity, including intrinsic donor and manufacturing process-driven heterogeneity (6), which together affect the potency attributes relevant to the clinical effectiveness of MSCs.

iPSCs differentiated into mesodermal derivatives (iMSCs) partially address this donor-intrinsic heterogeneity, deriving product from a single, clonal master cell bank rather than from repeated donor procurement (7, 8); however, parental differences (9) and batch-to-batch variability persist (10, 11). iMSCs recapitulate key functional properties of primary tissue-derived MSCs, including inducible indoleamine-2,3-dioxygenase (IDO) expression and inflammatory licensing responses (12). Their clonal origin and extended proliferative capacity also make them tractable to genetic engineering, allowing defined potency attributes to be modulated in a stable genetic background rather than selected for among donors. Despite these advantages, recent late-stage clinical trials of iMSC products have failed to meet their primary endpoints (13, 14), underscoring the urgent need to benchmark iMSC potency attributes mechanistically and functionally against primary tissue-derived MSCs.

Here, we report in-depth characterization of an immortalized, engineered iMSC cell line against multiple primary <u>a</u>dipose tissue-derived MSC(AT) donors. We used a curated panel of genes, anchored to functional assays, to define basal MSC(AT) fitness ranges for immunomodulatory and angiogenic functionalities (15). A Doxycycline-inducible, hTERT-immortalized, *TGFB1*-overexpressing iMSC line (16) was benchmarked against multiple primary MSC(AT) donors across morphometric, transcriptomic, miRNA, and functional dimensions. hTERT immortalization and doxycycline-inducible expression were used to sustain expansion capacity and to permit controlled transgene induction, respectively. In vivo, we evaluated the capacity of *TGFB1*-iMSCs to modulate the synovial joint macrophage compartment and joint pathology following intra-articular delivery in a murine destabilization of the medial meniscus (DMM) model of post-traumatic osteoarthritis (OA).

*TGFB1* was selected as the engineered payload because TGF-β protein correlated with immunomodulatory functional scores in our prior potency attribute analyses (15); higher *TGFB1* expression in bone <u>m</u>arrow MSC(<u>M</u>)s correlated with clinical efficacy in knee osteoarthritis (KOA) (17); and *TGFB1-*engineered cell lines are already under clinical investigation for KOA (18, 19, 20). These make *TGFB1* a rational candidate for augmenting immunomodulatory fitness in a defined genetic background for testing in KOA.

We found that *TGFB1*-iMSCs adopted a progenitor-like morphology and an angiogenic transcriptional signature, but *TGFB1* overexpression drove high immunomodulatory and moderate angiogenic functionality. In vivo, *TGFB1*-iMSC, but not primary tissue-derived MSC(AT), treatment altered the synovial macrophage phenotype without altering structural disease scores, supportive of its higher immunomodulatory capacity.

## Methods

### Generation and sourcing of iMSC cell lines

The *TGFB1*-overexpressing iMSC line was provided for research use by Pluristyx (Seattle, WA, USA). The line was developed by PanCELLa (Toronto, ON, Canada) using proprietary methods described in (16). Briefly, a single iPSC line derived from primary dermal fibroblasts was engineered to an MHC class I/II-null background through disruption of *B2M* and *CIITA*, followed by doxycycline-inducible immortalization via stable integration of an hTERT-2A-SV40 expression cassette. Immortalized iPSCs were differentiated to MSCs and further engineered by stable integration of a constitutively expressed human *TGFB1* transgene using the PiggyBac transposon system. All experiments in this study were performed using the resulting *TGFB1*-overexpressing derivative, hereafter termed *TGFB1-*iMSC.

iMSCs were expanded in StemXVivo Mesenchymal Stem Cell Expansion Media (R&D Systems, Minneapolis, MN, USA) at 37°C, 5% CO₂, under ambient air. Cells were passaged at approximately 80% confluency using TrypLE (Gibco, Waltham, MA, USA) and replated at 5,000 cells/cm². Prior to all assays, iMSCs were transitioned from StemXVivo to MesenCult™-ACF Plus medium (STEMCELL Technologies, Vancouver, BC, Canada) for 24 h, matching the medium used for primary MSC(AT) expansion and to benchmark against previous data (15). Primary MSC(AT) received an equivalent medium change in MesenCult™-ACF Plus over the same interval. Doxycycline (1 ug/mL; cat. D3072, Sigma-Aldrich, St. Louis, MO, USA) was maintained throughout cell expansion to sustain hTERT expression. At the transition to MesenCult-ACF Plus, doxycycline was either retained (+Dox) or withdrawn (-Dox) following three PBS washes, and this condition was maintained for the duration of each assay. All MSC experiments were performed between passage 3 and passage 5.

### Morphometric analysis

Single-cell suspensions were generated by TrypLE dissociation. Cell diameter and circularity were measured using the Vi-Cell XR Cell Counter (Beckman Coulter, Brea, CA, USA). Phase-contrast images were captured using an Evos XL Core at 10X objective.

For attachment efficiency, cells were plated at 5,000 cells/cm² and non-adherent cells were removed by washing at 4 h post-seeding. Adherent cells were harvested and counted, and attachment efficiency was expressed as a percentage of cells seeded. For proliferation, cells plated at 5,000 cells/cm² were harvested in log growth phase on days 3 and 5, and population doubling time was calculated over these 48 hours.

### Gene expression, microRNA profiling, and soluble factor measurements

Cells were plated in 24-well plates at 60,000 cells/well. Licensing was performed by addition of a cocktail of IFNγ (30 ng/mL), TNFα (10 ng/mL), and IL-1β (5 ng/mL) (Peprotech, Cranbury, NJ, USA) for 24 h. Unlicensed controls were cultured in parallel in the absence of pro-inflammatory cytokine stimuli. Following the 24 h licensing or unlicensed culture period, conditioned medium was collected, and cells were washed in PBS. Total RNA, including the small RNA fraction, was isolated from cells using the miRNeasy Mini Kit (Qiagen, Hilden, Germany) according to the manufacturer’s instructions. RNA concentration and purity were assessed on a DS-11 spectrophotometer (DeNovix, Wilmington, DE, USA), and RNA integrity was assessed using an Agilent 2100 Bioanalyzer (Agilent Technologies, Santa Clara, CA, USA); all samples had an RNA integrity number (RIN) > 9, and no samples were excluded on the basis of quality. The same RNA preparation was used as input for both mRNA and microRNA profiling.

Samples were run on the nCounter nSolver Analysis System (NanoString, Seattle, WA, USA). mRNA expression was assessed using a curated custom CodeSet, as previously reported (15); microRNA expression was assessed using the nCounter Human v3 miRNA Expression Assay (NanoString). Raw counts were imported into nSolver (v4.0) and assessed against the manufacturer’s quality control criteria for the Sprint Profiler (imaging, binding density, positive control linearity and limit of detection, and ligation controls). All samples passed and were retained. Counts were imported into R (v4.5.0). A per-sample detection threshold was defined as the mean negative control signal plus 2 SD; targets exceeding this threshold in at least two samples were retained (180/798 miRNAs; 16/17 mRNAs). Normalization and differential expression were performed for both datasets with NanoStringDiff (v.1.38.0) using glm.LRT, with Benjamini-Hochberg correction. Targets with adjusted p < 0.05 and |log₂ fold change| > 1 were considered differentially expressed. Volcano plots were generated with ggplot2 and ggrepel. Validated miRNA-gene interactions were retrieved using multiMiR (v1.30.0), and targets extracted separately for up-and downregulated miRNAs. Gene symbols were mapped to Entrez IDs with clusterProfiler (v4.16.0) and org.Hs.eg.db; unmapped genes were excluded. Gene Ontology and Reactome enrichment were performed using enrichGO and enrichPathway, with results shown as dot plots ordered by FDR-adjusted p-value. For integration, miRNA targets were filtered to differentially expressed genes from the mRNA dataset, classified by direction of regulation (Supp. Table 8), and exported for network visualization in Cytoscape.

Gene expression experiments in iMSCs and MSC(AT) were performed to i) pilot gene expression changes under doxycycline exposure and varying culture medium, and (ii) measure *TGFB1* and *TWIST1* expression. Both sets of experiments were performed under unlicensed and licensed conditions using the methods outlined above. RNA was isolated from iMSCs and MSC(AT)s by TRIzol-chloroform extraction, and cDNA was generated using SuperScript^TM^ IV VILO^TM^ Master Mix (Invitrogen, Waltham, USA). qPCR was run using custom primers (Suppl. Table 1, Invitrogen) and Taq Pro Universal SYBR qPCR Master Mix (Vazyme Biotech, Nanjing, China) on a QuantStudio^TM^ 5 system (ThermoFisher). Results were normalized (ΔΔCT) against reference genes (B2M, ACTB, and RPLP0) and presented as negative ΔΔCT values.

Conditioned medium samples were analyzed for soluble TGF-β1 concentrations by sandwich ELISA using the Human TGF-β1 DuoSet ELISA Development Kit (DY240, R&D Systems, Minneapolis, MN, USA) according to the manufacturer’s instructions.

### Monocyte/macrophage co-culture and phagocytosis

Peripheral blood-derived CD14⁺ monocytes were isolated from whole blood of healthy donors with informed consent (REB#14-7483) and cultured as previously described (15). Monocytes/macrophages were spiked with LPS (2.5 ng/mL; Millipore-Sigma) for 4 hours before collecting conditioned media for TNFα quantification by ELISA (R&D Systems, Minneapolis, MN, USA). Cells were harvested, and gene expression was measured by qPCR using a curated macrophage gene panel (Suppl. Table 2) as previously described (15).

Phagocytic activity was assessed in a separate experimental setup. Briefly, CD14⁺ monocytes were differentiated to macrophages over 5 days with 20 ng/mL M-CSF (PeproTech, USA) prior to 48hr treatment with MSC-conditioned medium. Phagocytosis of pHrodo™ Deep Red *E. coli* BioParticles® Conjugate (cat. P35360, ThermoFisher Scientific, Waltham, USA) was assessed according to the manufacturer’s protocol with a 1 h incubation, and fluorescence quantified by plate reader.

### HUVEC tube formation

Conditioned medium was prepared from iMSCs and MSC(AT)s, and the HUVEC tube formation assay was performed and analyzed by ImageJ Angiogenesis Analyzer, as previously described (15). EGM™-2 (Lonza, Basel, Switzerland) served as the positive control (POS) and basal medium without growth factors or supplements as the negative control (NEG). VEGF in conditioned medium was quantified by ELISA (R&D Systems, Minneapolis, MN, USA) according to the manufacturer’s instructions.

### Animal work

All animal procedures were approved by UHN Animal Research Center (AUP#5847). 15-16-wk-old, skeletally mature male C57BL/6 mice underwent destabilization of the medial meniscus (DMM) surgery as before (21, 22); healthy controls underwent no surgery. Three weeks post-DMM or no surgery, mice received a single intra-articular injection of 50,000 iMSCs or MSC(AT) in 5 µL of saline with 2.5% mouse serum, or vehicle alone (saline). Joints were harvested 8 weeks post-injection for flow cytometry, immunofluorescence, and histological analysis. 10 mice per group were used. Experiments were performed in batches to manage surgical burden, and normalization to DMM saline control groups was performed to account for batch differences.

For immunofluorescence, paraffin-embedded joint sections were deparaffinized in xylene, rehydrated through a graded ethanol series (100%, 95%, 70%, 50%), and subjected to antigen retrieval with Proteinase K Antigen Retrieval Solution (Abcam, ab64220) for 15 min at 37 °C. Sections were blocked in PBS containing 10% goat serum and 1% BSA for 1 h at room temperature, then incubated overnight at 4 °C with rat polyclonal anti-F4/80 (Bio-Rad, MCA497R, 1:100) together with either rabbit polyclonal anti-CD206 (Abcam, ab64693; 1:250) or rabbit polyclonal anti-MHC class II (Thermo Fisher Scientific, PA5-116876; 1:100). Sections were then incubated with goat anti-rat Alexa Fluor 647 (Invitrogen, A48265; 1:100) and goat anti-rabbit Alexa Fluor 488 (Invitrogen, A11034; 1:100) for 1 h at room temperature and mounted with DAPI-containing mounting medium (ab104139). Fluorescence images were acquired on a widefield Axio Observer microscope (Zeiss). Macrophage number (F4/80^+^ cells) was expressed per region of interest (ROI), and polarization was assessed as the ratio of MHCII to CD206 mean fluorescence intensity.

For single-cell flow cytometry, whole knee joints were dissected and enzymatically digested using collagenase Type I (250 U/mL; Sigma, C0130), hyaluronidase Type I-S (30 U/mL; Sigma, H3506), and DNase I Type II-S (30 U/mL; Sigma, D4513). Tissues underwent two 20-min digestion steps at 37°C with gentle agitation. Cell suspensions were sequentially filtered through 70-µm cell strainers, centrifuged at 300 × g for 5 min at 4°C, and resuspended in FACS buffer. Cells were stained with viability dye for 15 min at RT, followed by the flow cytometry antibody panel for 30 min at 4°C, and acquired on a BD SymphonyA3 cytometer (SickKids, Toronto, Canada). Data were analyzed using FlowJo. Macrophages were identified as CD64⁺MerTK⁺ cells, and the percentages and mean fluorescence intensities (MFI) of MHCII⁺ and CD206⁺ cells were quantified.

For histology, joints were fixed, paraffin-embedded, sectioned at 5 µm, and stained with toluidine blue. Cartilage degeneration was scored in the femur and tibia using the OARSI system (23), and synovitis and fibrosis were scored in the anterior and posterior compartments.

### Statistics

Plots were generated using GraphPad Prism (GraphPad Software, La Jolla, CA, USA). Unbiased hierarchical clustering (Ward method), principal component analysis, and desirability profiling were performed in JMP Pro. To resolve relative potency among conditions with comparable PC scores, Euclidean distance (ED) between conditions were calculated using PC1 and PC2 score values through 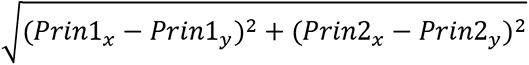, computed per-condition average to obtain between-group differences. All ED values were reported in Supplemental Table 8. One-way ANOVA with Tukey’s post-hoc test was used for comparisons across conditions, with groups sharing a letter considered not significantly different. Data are presented as mean ± SD unless otherwise indicated. Significance was defined as p < 0.05.

## Results

### Culture parameter selection and MSC(AT) donor benchmarking

Prior to comparing hTERT-immortalized, *TGFB1*-engineered iMSCs with primary MSC(AT) donors, culture parameters were first established to ensure appropriate benchmarking of cell phenotype and function. Candidate conditions were evaluated using a curated panel of established MSC(AT) potency attributes adapted from Robb et al. (15), together with proliferation analyses, under both unlicensed and licensed (pro-inflammatory cytokine stimulation) conditions. Based on these assessments, we selected a transition protocol in which iMSCs were cultured in StemXVivo and switched to MesenCult™-ACF Plus (TransitionM) for all subsequent experiments, matching the medium used for previous primary MSC(AT) expansion (Suppl. Fig. 1). To confirm that doxycycline (Dox) induction did not confound the measurements, iMSCs in TransitionM were assessed with and without Dox; no doxycycline-attributable differences were detected, and doxycycline exposure was therefore retained for subsequent experiments (Suppl. Fig. 2).

Four primary MSC(AT) donors were included. Three (AT1, AT3, and AT4) had previous benchmarking data (15), and one commercially sourced donor (AT2) was newly added and not previously characterized. Donors were re-designated AT1-AT4 here and do not correspond numerically to donor identifiers used previously. AT1 corresponded to the previous high-immunomodulatory potency donor while AT4 corresponded to the previously high-angiogenic potency donor; AT3 represented an intermediate profile. This design allowed all characterization readouts to be interpreted relative to MSC(AT) donors of known potency attributes.

### hTERT-immortalized, *TGFB1*-engineered iMSCs exhibited progenitor-like morphology with high proliferative capacity

Phase-contrast imaging demonstrated that hTERT-immortalized, *TGFB1*-engineered iMSCs retained the characteristic spindle-shaped morphology of primary tissue MSCs (Fig. 1A). Quantitative morphometric analysis demonstrated that iMSCs exhibited a significantly smaller cell diameter than primary MSC(AT), AT4, considered to be highly angiogenic, while remaining comparable to donors, AT1-AT3 (Fig. 1B). Consistent with this, iMSCs displayed significantly greater circularity than primary MSC(AT) donors, AT2-AT4 and was comparable to the highly immunomodulatory donor, AT1 (Fig. 1C). Assessment of attachment efficiency demonstrated that iMSCs attached comparably to donor AT2 and significantly more efficiently than donors AT1, AT3, and AT4 (Fig. 1D). Population doubling time was significantly shorter than primary MSC(AT) donor AT4 and comparable to other donors (Fig. 1E). Presence of Dox did not have any significant effect on morphology, attachment efficiency, or on doubling times (Suppl. Fig. 2A-D). These findings demonstrated that *TGFB1*-engineered iMSCs retained characteristic MSC morphology, albeit with smaller diameters and increased circularity, and their proliferative capacity was comparable to or greater than primary MSC(AT) donors.

**Figure 1.**
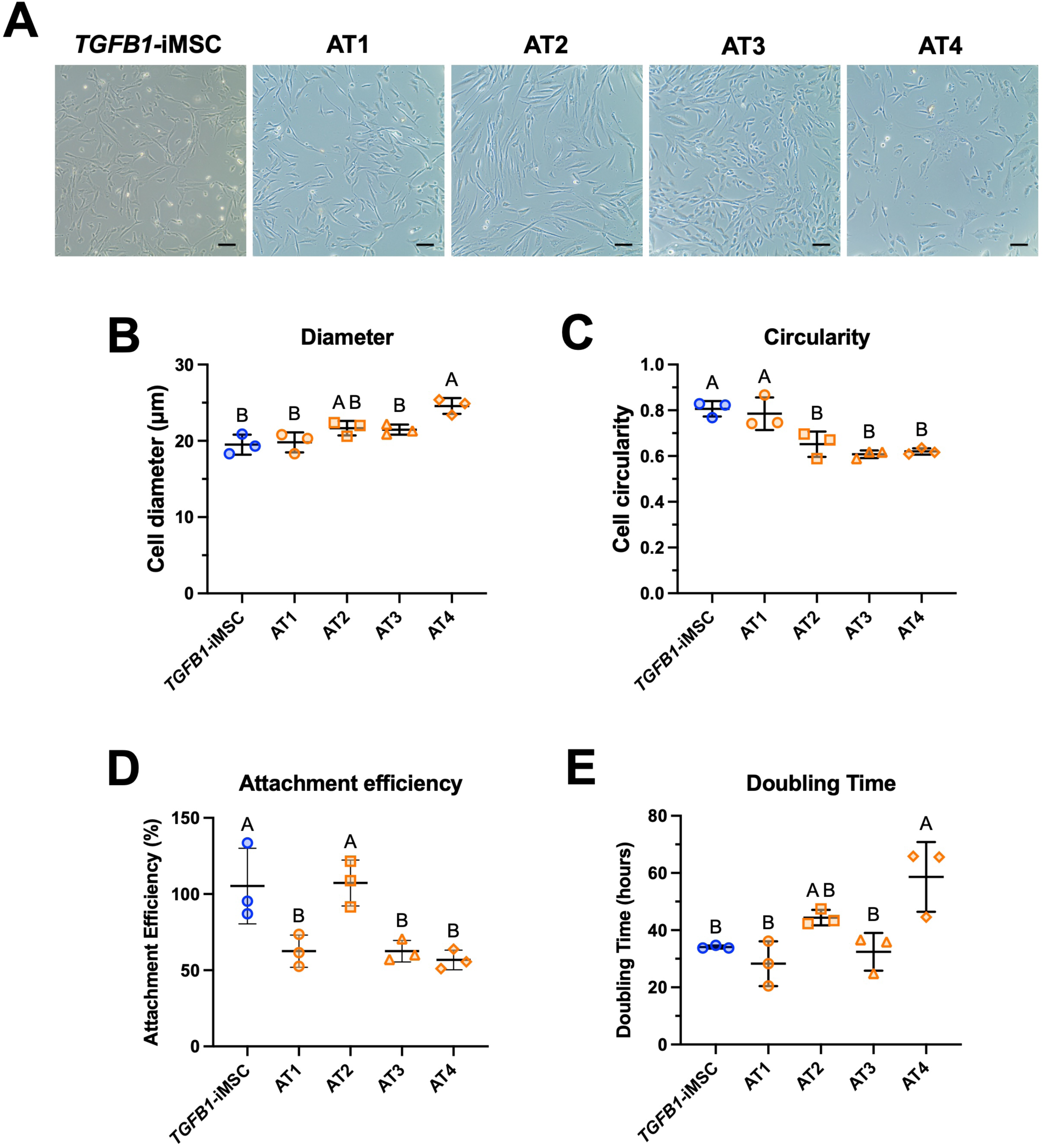
Morphometric and proliferative characterization of *TGFB1*-iMSCs and primary MSC(AT)s. (A) Representative phase-contrast images of *TGFB1-*iMSCs and four donor MSC(AT)s (AT1–AT4). All cells were used between passages 3 and 5. 10× magnification; scale bar = 50 μm. (B) Cell diameter (μm) and (C) cell circularity, measured by ViCell automated image analysis. (D) Attachment efficiency (%) quantified at 4 hours post-seeding. (E) Population doubling time (hours). Data are presented as mean ± SD, n = 3 independent experiments. Statistical comparisons were performed by one-way ANOVA with Tukey’s post hoc test; groups sharing a letter are not significantly different (p > 0.05).

### *TGFB1*-engineered iMSCs exhibited a distinct transcriptional identity independent of licensing status

To determine the transcriptional identity of *TGFB1*-engineered iMSCs, a curated panel of 17 immunomodulatory and angiogenic genes previously identified in MSC(AT) (15) and MSC(M) (24) were used. *TGFB1*-engineered iMSCs were benchmarked against previously characterized high-immunomodulatory AT1 and high-angiogenic AT4 donors (15). Unsupervised hierarchical clustering demonstrated that *TGFB1*-engineered iMSCs clustered separately from both MSC(AT) donors, which co-clustered together, under both licensed (Fig. 2A) and unlicensed (Fig. 2B) conditions. Relative to MSC(AT)s, *TGFB1*-engineered iMSCs expressed lower levels of *ANGPT1, ACTA2, HGF*, *CXCL8*, *VEGFA*, and *TNFAIP6*, and higher levels of *PDGFA*, *EDIL3*, *EDN1*, and *TSG101* regardless of licensing condition, demonstrating a transcriptomic profile of high immunosuppression. Dox exposure did not significantly affect gene expression of a short-listed panel, showing comparable expression levels of *TSG101*, *HGF*, *TNFAIP6*, and *PDGFA* (Suppl. Fig. 2E).

**Figure 2.**
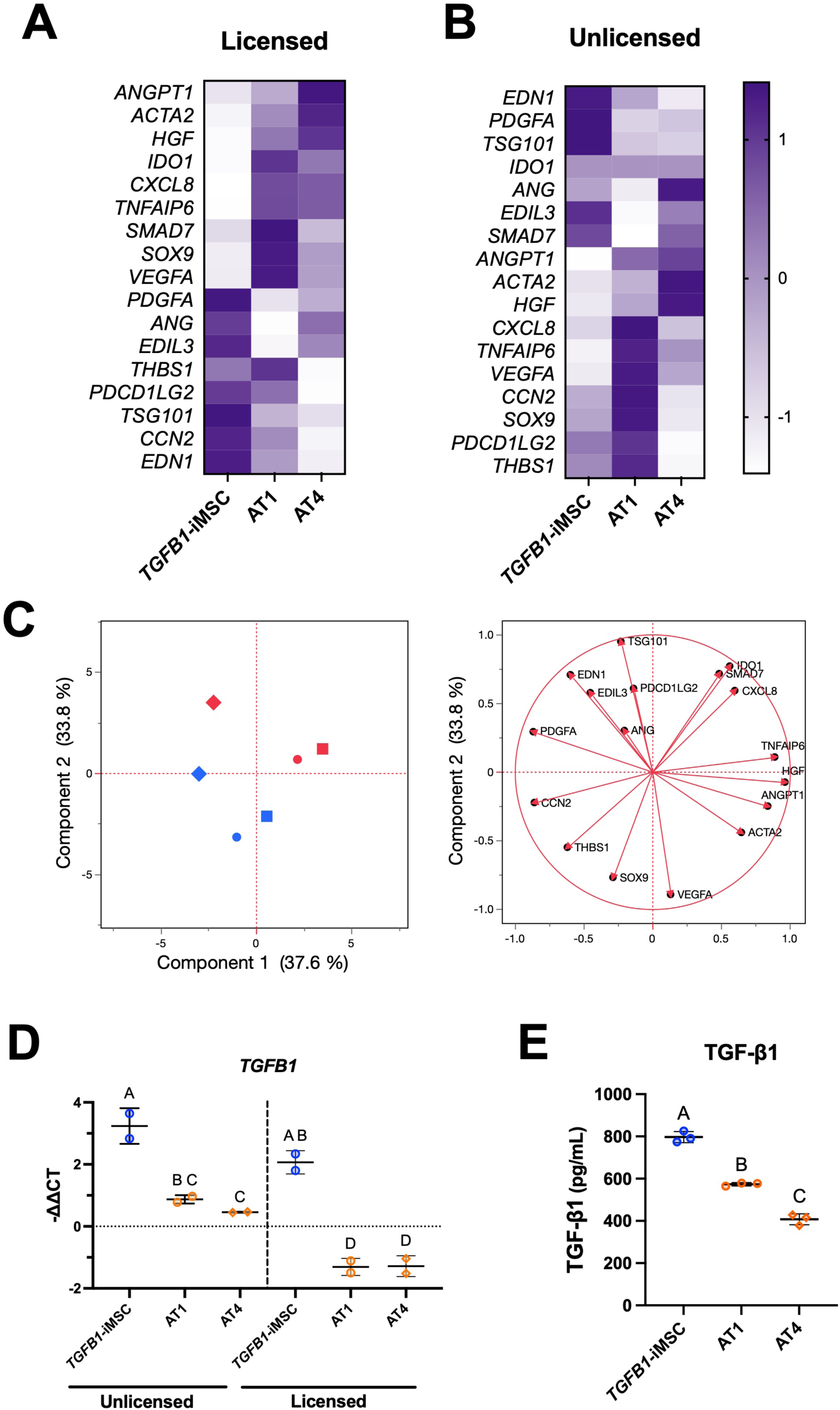
Gene expression profile of *TGFB1*-iMSCs under licensed and unlicensed conditions. NanoString nCounter gene expression profiling of a curated 17-gene immunomodulatory and angiogenic panel in *TGFB1*-iMSCs and in MSC(AT) from donors AT1 (high-immunomodulatory) and AT4 (high-angiogenic). Heatmaps of gene expression under (A) licensed and (B) unlicensed conditions, with genes ordered by unsupervised hierarchical clustering. (C) Principal component analysis (left) of all samples across both conditions, with the corresponding loading plot (right) showing each gene’s contribution to Component 1 and Component 2. Symbol colour denotes licensing condition (red, licensed; blue, unlicensed), and symbol shape denotes cell source (diamonds, *TGFB1-*iMSC; circles, AT1; squares, AT4). (D) *TGFB1* gene expression under unlicensed and licensed conditions, measured by RT-qPCR and shown as −ΔΔCT. (E) TGF-β1 protein secretion measured by ELISA in unlicensed conditioned medium. Data are mean ± SD (n = 2 or 3 per condition). Groups with different letters differ significantly (p < 0.05, one-way ANOVA with Tukey’s post hoc test).

Principal component analysis (PCA) stratified MSCs by tissue source and licensing status (Fig. 2C). PC1 (37.6% of variance) separated *TGFB1*-engineered iMSCs from MSC(AT) donors, which clustered together. PC2 (33.8%) separated licensed from unlicensed samples within each tissue source. Notably, the displacement between unlicensed and licensed states was similar in direction and magnitude for *TGFB1*-engineered iMSCs (ED = 3.612) and MSC(AT) donors (AT1-ED = 4.988; AT4-ED = 4.433), indicating that iMSCs behave like primary tissue MSC(AT) under licensing conditions despite distinct baseline transcriptional profiles. Loading analysis identified *HGF*, *TNFAIP6*, *PDGFA*, *CCN2*, and *ANGPT1* as the main contributors to PC1, whereas *TSG101*, *VEGFA*, *SOX9*, and *IDO1* contributed to PC2 (Suppl. Table 3).

*TGFB1* gene and protein expression were assessed, and as expected, *TGFB1*-engineered iMSCs exhibited elevated levels relative to primary MSC(AT) donors (Fig. 2D, E). Notably, inflammatory licensing suppressed *TGFB1* expression across *TGFB1*-engineered iMSCs and primary MSC(AT) groups, suggesting that licensing downregulates endogenous *TGFB1*. Licensing decreased *TWIST1* levels in *TGFB1-*iMSCs and donor AT1, but not in the high angiogenic donor, AT4 (Suppl. Fig. 3). As such, *TGFB1*-engineered iMSCs occupy a transcriptional state distinct from primary MSC(AT) donors, characterized by reduced expression of immunomodulatory mediators and enriched expression of matricellular and growth factor genes.

### A curated miRNA array showed *TGFB1*-engineered iMSCs clustering with high-performing MSC(AT) donors

To investigate whether post-transcriptional regulation contributes to the transcriptional differences observed between *TGFB1*-engineered iMSCs and MSC(AT) donors, we profiled the miRNA expression landscape of *TGFB1*-engineered iMSCs and MSC(AT) donors. PCA separated MSC(AT) donors along both PC1 (70.2%) and PC2 (16.6%), with *TGFB1*-engineered iMSCs occupying an intermediate position, close to both the high-immunomodulatory-ranked donor AT1 (ED = 1.886) and the high-angiogenic-ranked donor AT4 (ED = 3.564) (Fig. 3A). Differential expression analysis identified nine miRNAs significantly altered in *TGFB1*-engineered iMSCs relative to primary MSC(AT) (Fig. 3B). Gene targets included signaling by TGFβ family members and TGFβ receptor complex, alongside RHO GTPase cycling, growth factor receptor, and NTRK signaling (Fig. 3C, D). GO Biological Process enrichment converged on intracellular trafficking and localization in both directions, with macroautophagy and autophagy regulation specific to the downregulated set (Fig. 3E, F).

**Figure 3.**
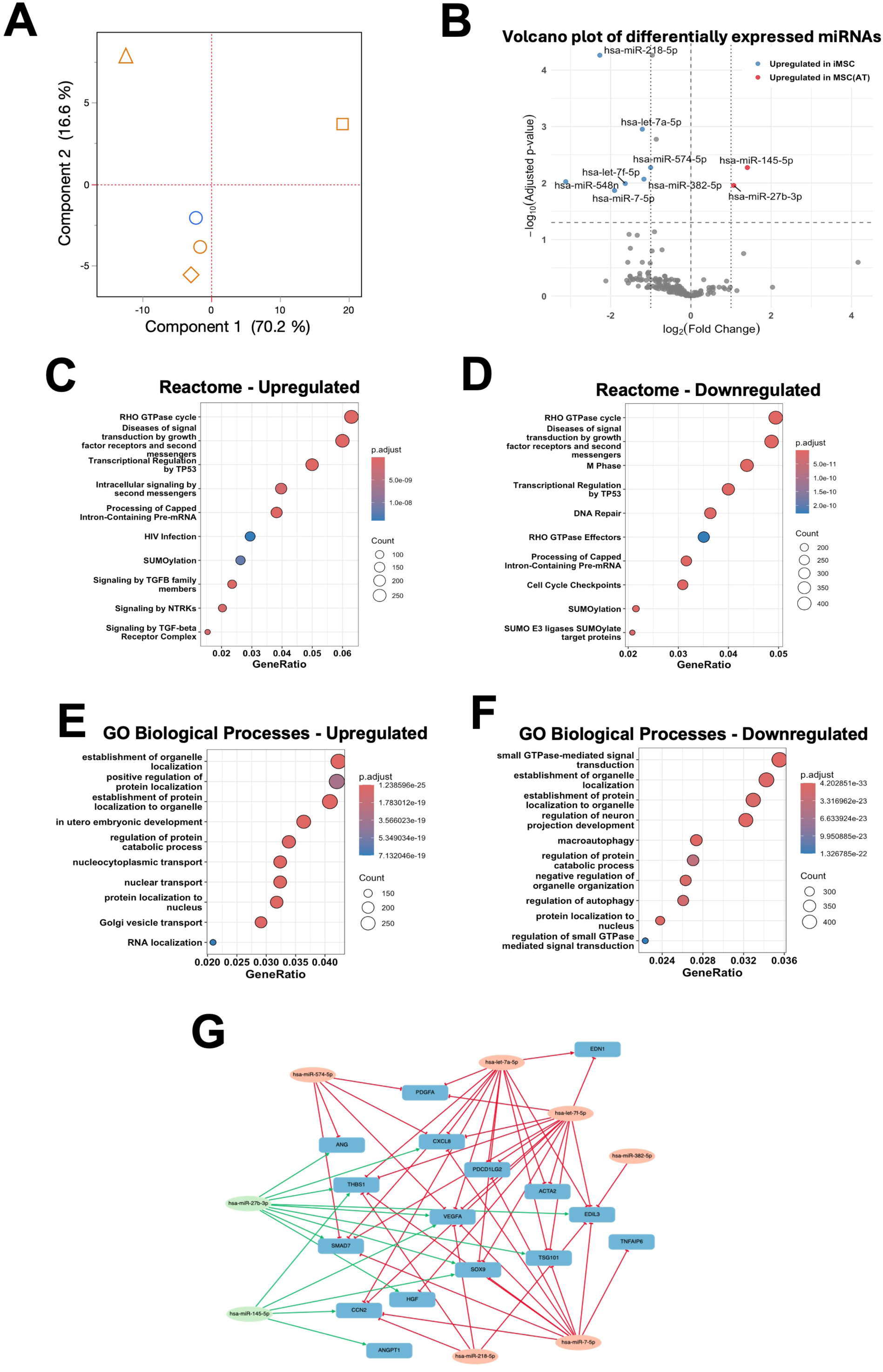
miRNA regulatory landscape clusters *TGFB1-*iMSCs with high-immunomodulatory and high-angiogenic MSC(AT) donors. (A) Principal component analysis of miRNA expression in iMSCs and MSC(AT) donors (AT1–AT4). Symbol colour denotes cell source (blue, *TGFB1-*iMSC; orange, MSC(AT) donors) and symbol shape denotes MSC(AT) donors (open circle, AT1; open square, AT2; open triangle, AT3; open diamond, AT4). (B) Volcano plot of differentially expressed miRNAs between *TGFB1-*iMSCs and MSC(AT) donors, showing log2(fold change) versus –log10(adjusted p-value). miRNAs upregulated in *TGFB1-*iMSCs are shown in blue; miRNAs upregulated in MSC(AT) are shown in red. (C, D) GO biological process enrichment of predicted gene targets of miRNAs upregulated (C) and downregulated (D) in iMSCs relative to MSC(AT). (E, F) Reactome pathway enrichment of predicted gene targets of miRNAs upregulated (E) and downregulated (F) in iMSCs relative to MSC(AT). Dot size represents gene count per pathway; colour represents adjusted p-value. (G) Network diagram of differentially expressed miRNAs and their predicted target genes, overlaid with the CQA gene panel used in Figures 2–4.

Predicted targets of these miRNAs were mapped against the subset of curated genes differentially expressed by iMSCs (Fig. 2), identifying six targeted genes (Fig. 3G). This relationship was directionally concordant for *ACTA2, TNFAIP6*, *CXCL8*, and *HGF*, which were reduced in *TGFB1*-engineered iMSCs alongside elevated let-7a-5p, let-7f-5p, miR-7-5p, and miR-574-5p. On the other hand, *EDIL3* and *TSG101* were elevated alongside reduced miR-27b-3p in *TGFB1*-engineered iMSCs. Predicted post-transcriptional targeting is therefore directionally consistent with the observed transcriptional differences across a portion of mapped genes. Together with the transcriptional profiling, these analyses establish that *TGFB1*-engineered iMSCs are molecularly distinct from primary MSC(AT) donors, congruent with their engineered and immortalized status. The top 20 upregulated and downregulated miRNAs in *TGFB1*-engineered iMSCs (Suppl. Table 4) and their predicted gene targets overlapping with tested mRNAs are provided (Suppl. Table 5).

### *TGFB1*-engineered iMSCs induced an intermediate pro-angiogenic response

We next evaluated the angiogenic functionality of *TGFB1*-engineered iMSCs. Vascular endothelial growth factor (VEGF) secretion was highest in the high-angiogenic potency-ranked donor AT4 and trended lower, albeit without significance, in the high-immunomodulatory potency-ranked donor AT1 (Fig. 4A). However, *TGFB1*-engineered iMSCs secreted significantly less VEGF than the MSC(AT) angiogenic donor, AT4.

**Figure 4.**
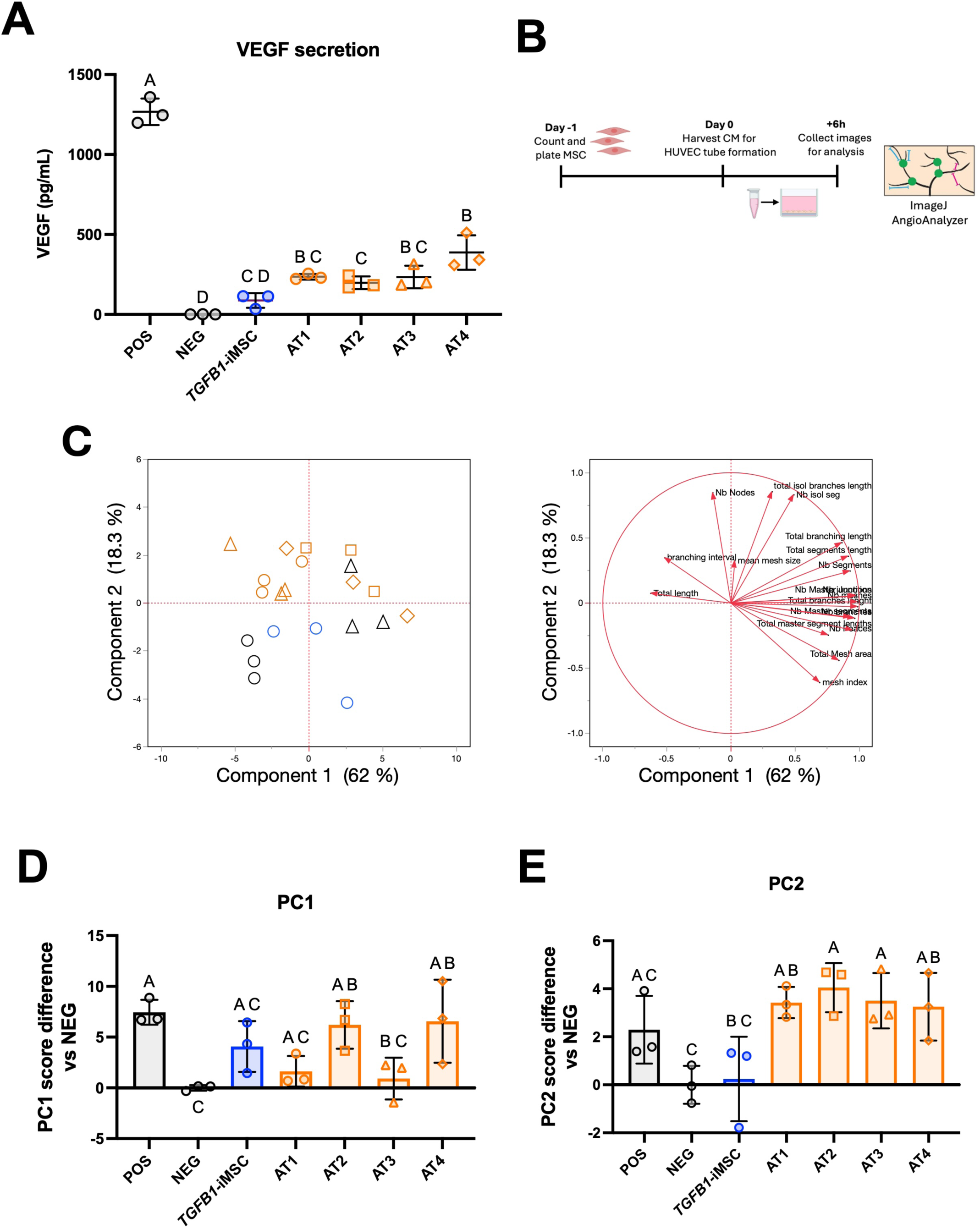
*TGFB1-*iMSCs drive an intermediate pro-angiogenic functional response in HUVECs. (A) VEGF secretion (pg/mL) from *TGFB1-*iMSC- and MSC(AT)-conditioned media, measured by ELISA. (B) Schematic of the HUVEC tube formation assay. MSCs were counted and plated; conditioned medium (CM) was harvested and applied to HUVECs, and tube formation images were collected 6 hours later for analysis by ImageJ AngioAnalyzer. C) Principal component analysis of HUVEC tube formation parameters following treatment with *TGFB1-*iMSC- or MSC(AT)-conditioned medium, positive control (POS), or negative control (NEG) (left), with corresponding loading plot showing tube formation parameter contributions to Component 1 and Component 2 (right). Symbol colour denotes cell source (black, controls; blue, *TGFB1-*iMSC; orange, MSC(AT) donors), and symbol shape denotes controls (open circle, negative; open triangle, positive) and MSC(AT) donors (open circle, AT1; open square, AT2; open triangle, AT3; open diamond, AT4). (D) PC1 and (E) PC2 score differences relative to NEG. Data are presented as mean ± SD, n = 3 biological replicates. Statistical comparisons were performed by one-way ANOVA with Tukey’s post hoc test; groups sharing a letter are not significantly different (p > 0.05).

Using human umbilical vein endothelial cells (HUVECs), we performed vessel network formation, as previously described (Fig. 4B) (15), with representative vessel images across treatment groups shown (Suppl. Fig. 4). PCA resolved 80.3% of total variance across the first two components (Fig. 4C). Increased total branch length, number of meshes, and number of master junctions were the main contributors to higher scores along the PC1 axis (62.0% of variance), indicative of greater angiogenesis (Suppl. Table 6). A greater total isolated branch length, number of nodes, and number of isolated segments drove higher scores along the PC2 axis (18.3% of variance) (Suppl. Table 6). In the PCA plot, HUVECs treated with *TGFB1*-engineered iMSC-conditioned medium were near-equidistant from all other conditions tested, separating from the negative (ED = 4.084) and positive (ED = 3.947) controls and falling between the primary MSC(AT) donors (AT1-ED = 4.012; AT2-ED = 4.359; AT3-ED = 4.535; AT4-ED = 3.905) (Suppl. Table 8). This pattern reflects an intermediate iMSC position along PC1 combined with displacement from all other MSC(AT) conditions along PC2. Consistent with this, PC1 scores did not differ significantly between groups, with *TGFB1*-engineered iMSCs trending intermediate (Fig. 4D). On PC2, *TGFB1*-engineered iMSCs differed from AT2 and AT3 and, unlike all four primary donors, were indistinguishable from the negative control (Fig. 4E). Presence of Dox did not alter the intermediate angiogenic functionality of *TGFB1-*engineered iMSCs (Suppl. Fig. 2F-G). Collectively, *TGFB1*-engineered iMSCs induced an intermediate pro-angiogenic functional response, relative to primary MSC(AT) donors, but with a distinct network architecture profile.

### *TGFB1*-engineered iMSCs exhibited equivalent or greater immunomodulatory potency than primary MSC(AT)s

We next assessed whether this intermediate angiogenic functionality was mirrored by immunomodulatory functionality. Peripheral blood-derived CD14^+^ monocytes were differentiated into macrophages and co-cultured with *TGFB1*-engineered iMSCs or primary MSC(AT) donors, as previously described (Fig. 5A) (15). Macrophages co-cultured with *TGFB1*-engineered iMSCs secreted significantly less TNFα than untreated controls and primary donors AT2-AT4; TNFα was also non-significantly lower compared to the high immunomodulatory-potency donor AT1 (Fig. 5B). Congruent with this, the phagocytic activity of *TGFB1*-engineered iMSC-treated macrophages was significantly better than that of the high-angiogenic-ranked MSC(AT) donor, AT4 and comparable to the high-immunomodulatory donor, AT1 (Fig. 5C). Presence of Dox did not significantly induce additional immunosuppression in the co-culture gene expression nor TNFα production (Suppl. Fig. 2H-I).

**Figure 5.**
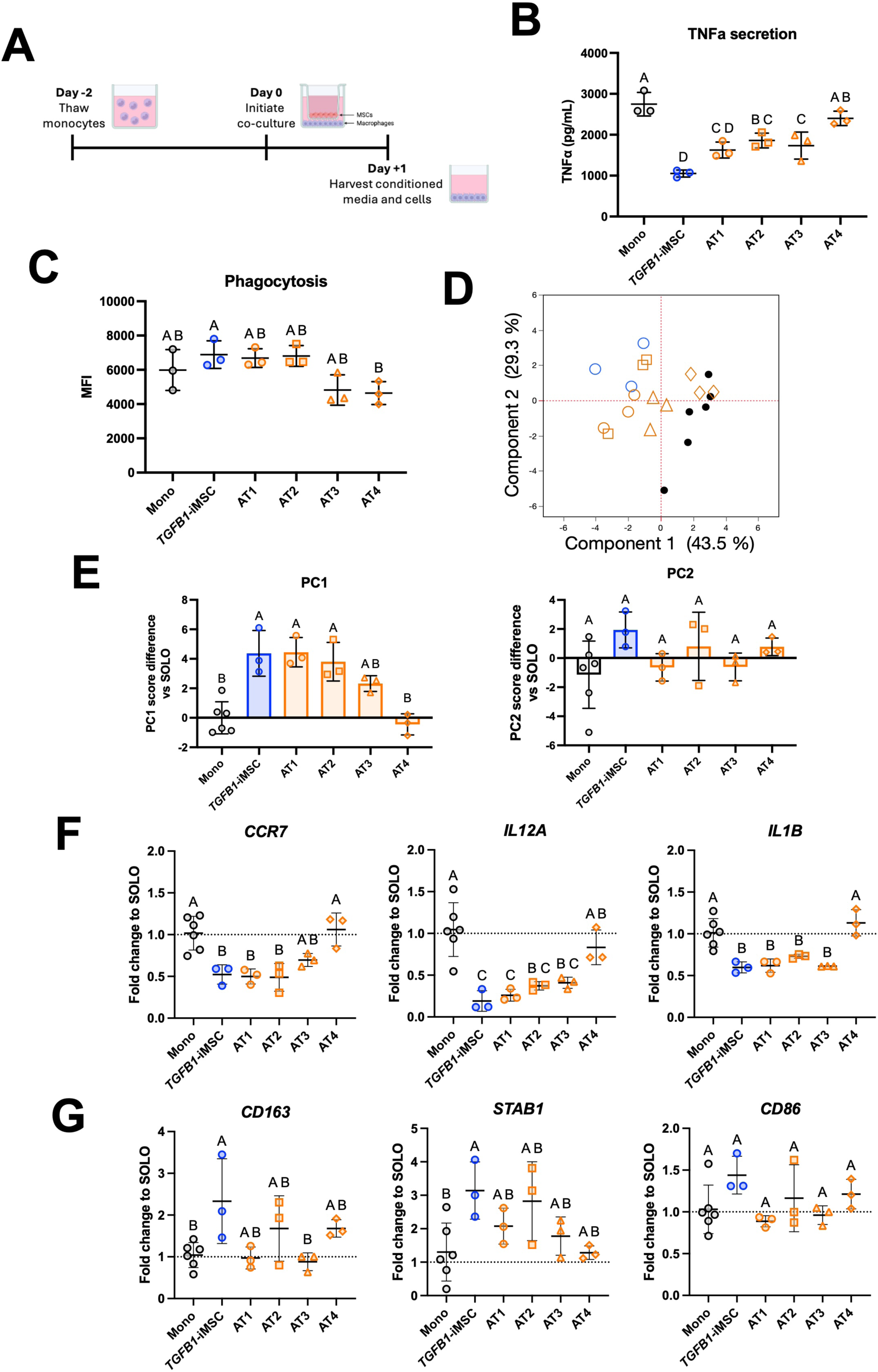
*TGFB1-*iMSCs exhibit equal or higher immunomodulatory potency relative to primary MSC(AT) donors. (A) Schematic of the monocyte/macrophage co-culture experimental workflow. CD14+ monocytes were differentiated and co-cultured with *TGFB1-*iMSCs or MSC(AT) donors (AT1-AT4) in transwell inserts for 24 hours, followed by harvest of conditioned medium and cells. (B) TNFα secretion (pg/mL) measured by ELISA. (C) Phagocytic activity measured by mean fluorescence intensity (MFI). (D) Principal component analysis of monocyte/macrophage gene expression following co-culture with *TGFB1-*iMSC or MSC(AT) donors. Symbol colour denotes cell source (black, monocyte control; blue, *TGFB1-*iMSC; orange, MSC(AT) donors) and symbol shape denotes MSC(AT) donors (open circle, AT1; open square, AT2; open triangle, AT3; open diamond, AT4). (E) PC1 (left) and PC2 (right) score differences relative to Mono (monocyte-only control). (F) Fold change relative to Mono for CCR7, IL12A, and IL1B, markers of pro-inflammatory (M1-like) polarization. (G) Fold change relative to Mono for CD163, STAB1, and CD86, markers of anti-inflammatory (M2-like) polarization. Data are presented as mean ± SD, n = 3 independent experiments. Statistical comparisons were performed by one-way ANOVA with Tukey’s post hoc test; groups sharing a letter are not significantly different (p > 0.05).

PCA of curated macrophage gene panel showed that macrophages co-cultured with *TGFB1*-engineered iMSCs clustered with high-immunomodulatory MSC(AT) donors AT1-AT3 (AT1-ED = 2.575; AT2-ED = 1.264; AT3-ED = 3.267), while being furthest from the high-angiogenic potency-ranked donor, AT4 (ED = 4.952) and macrophage-alone controls (ED = 5.345) (Fig. 5D; Suppl. Table 8). The PC1 axis (43.5% of variance) was driven primarily by the pro-inflammatory genes *CCR7*, *IL12A*, and *IL1B* (Fig. 5E; Suppl. Table 7), all of which were reduced by co-culture with *TGFB1*-engineered iMSCs and with donors, AT1-AT3, but not with the high angiogenic potency ranked donor AT4 (Fig. 3F). The PC2 axis (29.3% of variance) was driven primarily by the anti-inflammatory genes *CD163*, *STAB1*, and *CD86* but PC2 scores did not differ significantly across groups (Fig. 5E, F). A full analysis of a broader panel of macrophage polarization genes similarly demonstrated consistent immunomodulatory effects following *TGFB1*-engineered iMSC co-culture (Suppl. Fig. 5). Together, these findings demonstrate that *TGFB1*-engineered iMSCs exhibit immunomodulatory activity comparable to or exceeding that of the high-immunomodulatory ranked primary MSC(AT) donor. Notably, this effect was not mediated by Dox, given that *TGFB1-*engineered iMSCs maintained this effect in the absence of Dox (Suppl. Fig. 2H-I).

### *TGFB1*-engineered iMSCs reduced synovial macrophage number and promoted anti-inflammatory macrophage population

To evaluate whether the in vitro immunomodulatory potency of *TGFB1*-engineered iMSCs translated to a complex disease setting, we used a chronic model of mechanical instability-induced osteoarthritis. Skeletally mature mice underwent destabilization of medial meniscus (DMM) surgery at 15 weeks of age, followed by intra-articular injection of either *TGFB1*-engineered iMSCs, high-immunomodulatory AT1, or saline, 3 weeks post-DMM, with joints harvested 8 weeks post-injection for longer-term assessment of MSC effects (Fig. 6A).

**Figure 6:**
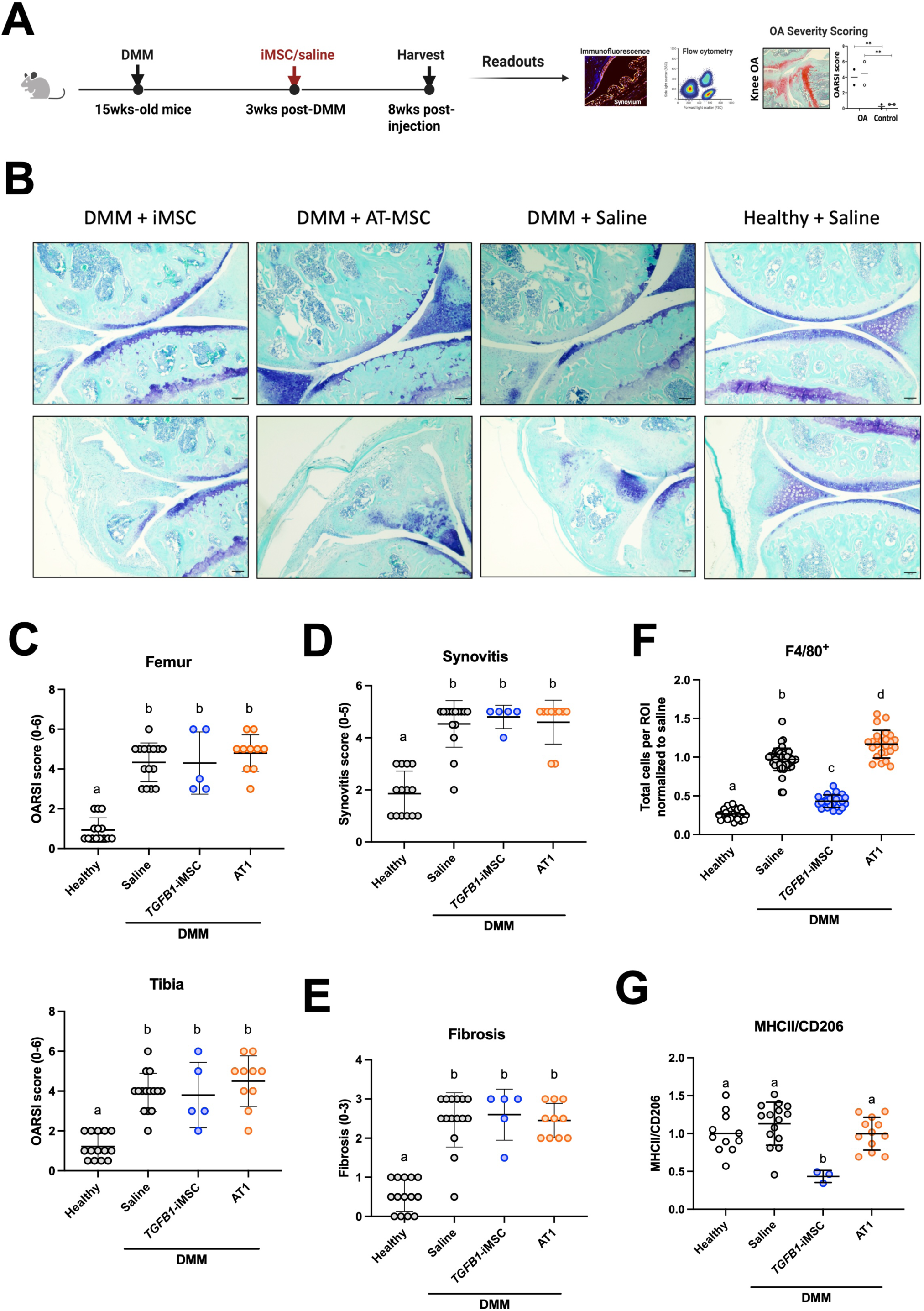
*TGFB1-*iMSCs reduced the total number of synovial macrophages and decreased the ratio of pro-inflammatory marker-expressing macrophages. (A) Experimental timeline. Destabilization of the medial meniscus (DMM) surgery was performed on 15-week-old mice. At 3 weeks post-DMM, mice received an intra-articular injection of *TGFB1-*iMSCs, high-immunomodulatory primary MSC(AT) donor, AT1, or saline. Knee joints were harvested 8 weeks post-injection for immunofluorescence, flow cytometry, and OA severity scoring. (B) Representative Toluidine Blue-stained sagittal sections of knee joints from DMM + *TGFB1-*iMSC, DMM + AT1, DMM + saline, and healthy + saline mice. Top row: articular cartilage images used for OARSI scoring. Bottom row: synovium images used for synovitis and fibrosis scoring. Scale bar = 100 µm. (C) OARSI scores (0–6) of the femur (left) and tibia (right). (D) Synovitis scores (0 - 6). (E) Synovial fibrosis scores (0 - 3). (F) Total number of F4/80⁺ synovial macrophages per region of interest (ROI), normalized to DMM + saline. (G) Ratio of MHCII⁺ (pro-inflammatory) to CD206⁺ (anti-inflammatory) synovial macrophages. Data are mean ± SD, n= 10 mice/group. Groups with different letters differ significantly (p < 0.05).

Histological assessment (Fig. 6B) showed no difference in cartilage degeneration between *TGFB1*-engineered iMSC, primary MSC(AT) or saline groups, with all groups displaying comparably elevated standardized cartilage degradation scores (23) in the femur and tibia relative to healthy controls (Fig. 6C). Synovitis (Fig. 6D) and fibrosis (Fig. 6E) scores were similarly indistinguishable between all three groups.

Immunofluorescence analysis revealed that saline and primary MSC(AT), donor AT1-treated mice exhibited a significant increase in total F4/80⁺ macrophages per region of interest (ROI) relative to healthy controls; relative to saline controls, F4/80+ macrophages were significantly reduced in *TGFB1*-engineered iMSC-treated mice (Fig. 6F). The major histocompatibility complex class II (MHCII)/CD206 mean fluorescence intensity ratio, reflecting the phenotypic balance of pro- to anti-inflammatory macrophages, did not differ significantly between healthy control and saline groups, but were significantly reduced in *TGFB1*-engineered iMSC-treated compared to saline controls and primary MSC(AT)-treated mice (Fig. 6G). Data are reported relative to matched saline controls to account for batch differences. Together, these data indicate that *TGFB1*-engineered iMSC-treated mice showed differential macrophage modulation compared with primary MSC(AT)-treated mice, without concomitant changes in cartilage, synovial inflammation, or fibrosis at the pre-selected endpoint.

## Discussion

This study established a doxycycline-inducible, hTERT-immortalized, *TGFB1*-overexpressing iMSC line as a benchmarked research tool for interrogating MSC potency attributes and applies a previously established potency framework to position it relative to primary MSC(AT) donors. *TGFB1*-engineered iMSCs displayed a progenitor-like morphology, a transcriptional profile weighted toward matricellular and growth factor genes, a distinct miRNA profile, intermediate in vitro angiogenic functionality, and high immunomodulatory potency. In a chronic post-traumatic osteoarthritis model, *TGFB1*-engineered iMSCs were comparable to primary MSC(AT) in structural and inflammatory outcomes. However, *TGFB1*-engineered iMSCs, but not MSC(AT), reduced total synovial macrophage numbers and promoted a higher ratio of anti-inflammatory to pro-inflammatory macrophages, relative to matched controls, an effect that persisted to eight weeks. Together, these findings demonstrate that iMSCs can be engineered to have desirable attributes compared to primary MSCs that may be fit-for-purpose for a given indication.

While iMSCs are a promising tool for largely eliminating donor-to-donor variability, donor-independent manufacturing does not by itself guarantee consistency or efficacy. For example, iMSC-derived EV potency still varied across manufacturing batches (10) and with passage number (11), and notably, early reports show that clinical outcomes in complex indications, including graft-versus-host disease and knee osteoarthritis, have been unpromising (13, 14).

Variability is therefore displaced from the donor to the manufacturing process, where it becomes more tractable to cell engineering or tighter control of process parameters.

Furthermore, immortalized iMSC lines offer improved expansion potential and a scalable source of MSC-derived extracellular vesicles, making them useful research tools for comparing mechanistic attributes across centres and disease indications (25, 26, 27). That advantage, however, is bounded by clonal selection and epigenetic drift during extended passaging, which can alter cell behaviour and reintroduce heterogeneity (28, 29, 30). Cell lines must therefore be used at early passages, stringently controlled, and matched across batches. Within these limits, the line’s scalability and amenability to genetic modification make it a mechanistically informative tool for dissecting the determinants of engineered MSC potency.

Consistent with earlier studies (15, 24), our primary MSC(AT) donors reproduced the previously established hierarchy in which highly angiogenic donors exhibit comparatively lower immunomodulatory potency, and vice versa. A similar relationship has been described in bone marrow MSCs along the TWIST1-TSG6 expression axis (31, 32, 33) and was observed here: the high-immunomodulatory donor AT1 had low *TWIST1* and high *TNFAIP6* expression, while the high-angiogenic donor AT4 showed the inverse. The immortalized *TGFB1*-engineered iMSCs did not follow this pattern. Despite low *TNFAIP6* and moderate *TWIST1* expression, and a transcriptional signature more angiogenically weighted than that of AT4, they displayed only moderate angiogenic activity and high immunomodulatory activity compared to the high-immunomodulatory AT1. The absence of the expected *TWIST1/TNFAIP6* relationship suggests that *TGFB1* overexpression supplies an alternative, TSG-6-independent route to immunomodulatory activity. TGF-β1 is a direct driver of macrophage polarization towards an anti-inflammatory phenotype and suppresses macrophage TNF-α and nitric oxide production (34), and MSC-secreted TGF-β1 has likewise been implicated in the same process (35, 36). Further, our functional data support this: our MSC(AT) high angiogenic donor, AT4, and high-immunomodulatory donor, AT1, displayed high vs. low vessel connectivity, respectively, aligning with previous data (15), whereas *TGFB1*-engineered iMSCs displayed intermediate connectivity, falling between the two donor extremes. Thus, our data support that *TGFB1*-engineered iMSCs have high immunomodulatory functionality, despite a high angiogenic transcriptomic signature.

Post-transcriptional regulation appears to contribute to this reconfiguration. Differentially expressed miRNAs and their predicted targets were directionally concordant across all mapped genes, with elevated let-7a-5p, let-7f-5p, miR-7-5p, and miR-574-5p accompanying reduced *ACTA2*, *TNFAIP6*, *CXCL8*, and *HGF*, respectively, and reduced miR-27b-3p accompanying elevated *EDIL3* and *TSG101*. Enrichment of TGFβ family and TGFβ receptor complex signalling terms among the targets of iMSC-upregulated miRNAs provides additional post-transcriptional support. The pathways enriched here include cell cycle (GTPases), TGFβ signalling, and NTRK signalling. Interestingly, many of these pathways were also enriched in miRNA profiling of MSC(M)s, classified as function-pain responders (24) in our KOA clinical trial (17). This implies that *TGFB1* expression and signalling contributed to the functional improvements we previously observed with MSC(M) in KOA (17).

In a complex, chronic post-traumatic osteoarthritis (OA) model, a single intra-articular *TGFB1*-engineered iMSC injection reduced total synovial macrophage numbers and increased the ratio of macrophages expressing anti-inflammatory-to-pro-inflammatory markers, relative to controls; this reduction or switch in total macrophages was not replicated by the single high immunomodulatory primary MSC(AT) donor, AT1. We ascribe this effect to an engineered *TGFB1* input, consistent with its established action on the myeloid compartment (35, 36).

*TGFB1* has clinical precedent as a driver of macrophage-directed activity in the joint. A *TGFB1*-overexpressing cell therapy, although missing both co-primary and all secondary endpoints in a Phase III trial (37), performed well in two prior smaller Phase II trials (19, 38), acting primarily through synovial macrophage polarization. These anti-inflammatory effects were also attributed to *TGFB1*-induced prostaglandin E2 production (20). Our findings extend this precedent to an engineered iMSC platform, indicating that a selective gene editing strategy can enhance macrophage-modulating function intrinsic to MSC biology. Importantly, this genetic enhancement was not accompanied by increased joint fibrosis in our post-traumatic OA model, positioning *TGFB1* overexpression as a rational and safe approach to engineering MSCs toward more potent immunomodulation.

The effects on synovial macrophages persisted to eight weeks, exceeding the four-week residence we previously reported for isogenic bone marrow MSCs in a post-traumatic OA joint (22), suggestive of durable reprogramming of the synovial macrophage compartment, potentially through stable transcriptional, epigenetic, or other regulatory mechanisms that remain active after cell clearance. MSCs are known to durably reprogram immune output through central mechanisms acting on bone marrow progenitors, leaving hematopoietic stem cells poised towards enhanced myelopoiesis months after exposure (39). While intra-articular injection is unlikely to reach the bone marrow compartment at a sufficient dose, emerging evidence suggests that MSCs can also directly imprint peripheral immune cells (40, 41). Two relevant, ontogenetically distinct synovial macrophage populations offer plausible targets for such imprinting: embryonically derived, self-renewing tissue-resident macrophages that form a stable lining-layer barrier maintained independently of circulating precursors, as we (42) and others (43) have shown, and monocyte-derived macrophages recruited during joint injury that persist as long-lived synovial residents (42, 44). Either could sustain a stable imprint initiated during the brief window of iMSC exposure through previously reported mechanisms (45, 46).

The changes in synovial macrophages were not accompanied by improvements in synovitis, fibrosis, or cartilage degradation, with either *TGFB1*-engineered iMSCs or MSC(AT)s, echoing recent top-level data from non-engineered iMSCs in a KOA clinical trial (14). The negative results may also reflect our choice of a post-traumatic OA model, driven by mechanical instability with comparatively mild inflammation (47, 48). Previous studies using unmodified murine MSC(M)s have similarly failed to improve cartilage degradation, synovitis, and fibrosis scores in DMM (22), including the control unengineered compact bone-derived MSCs (49). Notably, in that study, the engineered IL-1β sticky-trap MSC line did significantly attenuate cartilage degradation and synovitis, suggesting that structural benefit in DMM may require direct interception of a cartilage catabolic driver rather than immunomodulation alone (49). More inflammatory models, albeit non-pathologically relevant, such as collagenase-induced OA (CIOA) (48, 50), have shown more robust MSC-mediated reductions in synovial inflammation and cartilage degradation ( 51). Given evidence that MSCs may be more effective in a subset of patients with more inflammatory OA (24, 52), the DMM model provides a limited window for MSCs to demonstrate their full therapeutic potential.

Another limitation of our approach is that the use of curated molecular and post-transcriptional characterization of *TGFB1*-engineered iMSCs to primary MSC(AT) was limited to a single iMSC replicate, precluding formal statistical testing. Expanding this to several *TGFB1*-engineered iMSC batches, and to full single-cell-based sequencing would generate batch-to-batch variability and unbiased datasets, useful for querying full transcriptomic and miRNA changes. miRNA-target relationships reported here are computationally predicted rather than experimentally validated, and their directional concordance is based on correlation. The in vivo study examined a single treatment regimen, dose, and endpoint, precluding assessment of temporal changes in macrophage reprogramming and other tissue repair readouts.

Collectively, this study demonstrates that an hTERT-immortalized, *TGFB1*-engineered iMSC line demonstrated equivalent or higher immunomodulatory potency in vitro and in vivo, compared to primary MSC(AT) donors. These data serve as proof-of-concept for engineering iMSCs to enhance specific attributes for a given indication. The use of immortalized iMSCs may also serve as a useful research tool to reduce donor-to-donor and batch-to-batch variability, especially for larger, consortium-based studies as suggested (53). Together, this *TGFB1*-engineered iMSC cell line served as a tractable platform for studying the molecular and functional consequences of engineered MSCs.

## Abbreviations

MSCs: Mesenchymal stromal cells
iPSCs: Induced pluripotent stem cells
iMSCs: Induced pluripotent stem cell-derived mesenchymal stromal cells
MSC(AT)s: Adipose tissue-derived mesenchymal stromal cells
MSC(M)s: Bone marrow mesenchymal stromal cells
KOA: Knee osteoarthritis
DMM: Destabilization of medial meniscus
FDA: Food and Drug Administration IDO: Indoleamine-2,3-dioxygenase Dox: Doxycycline
hTERT: Human telomerase reverse transcriptase
LPS: Lipopolysaccharide
HUVECs: human umbilical vein endothelial cells
TGFB1: Transforming growth factor beta-1
VEGF: Vascular endothelial growth factor
TNFα: Tumour necrosis factor alpha
ED: Euclidean distance

## Acknowledgments

This work is supported by Eterna Health. The work is also supported by the Schroeder Arthritis Institute via the University Health Network Foundation. Schematics were developed using BioRender.com.

## Author Contributions

K.F. performed experiments, conducted all statistical analyses, generated figures, and drafted and revised the manuscript. M.R. performed experiments. R.K. performed statistical analysis. J.G., R.L., A.Z., S.N., and S.S. performed animal experiments and statistical analysis. K.R. conceived the study and contributed to manuscript editing. R.G. secured funding and contributed to manuscript editing. S.V. conceived and supervised the study, and contributed to writing and editing the manuscript. All authors reviewed and approved the final manuscript.

