## Supplemental Tables for "*TGFB1*-Engineered Induced Mesenchymal Stromal Cells Exhibit High Immunomodulatory Potency and Durably Reprogram Synovial Macrophages in Osteoarthritis"

**Supplemental Table 1. Mesenchymal stromal cell donor information**

| Donor ID | Sex | Age | Depot |
| --- | --- | --- | --- |
| iPSC (iMSC) | N/A | N/A | Skin fibroblasts |
| AT2 | F | 47 | Commercial Product |

**Supplemental Table 2. Primer sequences**

| Gene | Panel | Forward | Reverse |
| --- | --- | --- | --- |
| *B2M* | Housekeeping | CTCCGTGGCCTTAGCTGTG | TTTGGAGTACGCTGGATAGCCT |
| *ACTB* | Housekeeping | AGAGGGAAATCGTGCGTGAC | AGGAGCCAGGGCAGTAATC |
| *RPL13A* | Housekeeping | TGCGACAAAACCTCCTCCTT | TGTTGATGCCTTCACAGCGTA |
| *TGFB1* | MSC | CTAATGGTGGAAACCCACAACG | TATCGCCAGGAATTGTTGCTG |
| *TWIST1* | MSC | TCCATTTTCTCCTTCTCTGGAA | CCTTCTCGGTCTGGAGGAT |
| *CCR7* | Monocyte | TTTTACCGCCCAGAGAGCG | AATGACAAGGAGAGCCACC |
| *IL12A* | Monocyte | CCTCCACTGTGCTGGTTTTAT | TCAGCAACATGCTCCAGAAG |
| *IL1B* | Monocyte | GTACCTGTCCTGCGTGTTGA | GGGAACTGGGCAGACTCAAA |
| *CD163* | Monocyte | TGGACCTAATGAATTCCTCAGAAAA | ACACAGAAATTAGTTCAGCAGCA |
| *CD206* | Monocyte | CTACAAGGGATCGGGTTTATGGA | TTGGCATTGCCTAGTAGCGTA |
| *STAB1* | Monocyte | GAACCATGTGCCACTGGAAGGC | AGCGGAATCTCCTGGTGCAGTT |
| *CD86* | Monocyte | CTGCTCATCTATACACGGTTACC | GGAAACGTCGTACAGTTCTGTG |
| *TREM1* | Monocyte | AGTTGCAGCTCGGAGTTCTGAGACA | GAACCATGTGCCACTGGAAGGC |
| *IL10* | Monocyte | CGAGATGCCTTCAGCAGAGT | CGCCTTGATGTCTGGGTCTT |
| *HLA-DRA* | Monocyte | AAGCACTGGGAGTTTGATGC | ATTGCTTTTGCGCAATCCCT |
| *PDL1* | Monocyte | GCTGCACTAATTGTCTATTGGGA | AATTCGCTTGTAGTCGGCACC |
| *HMOX1* | Monocyte | AAGACTGCGTTCCTGCTCAAC | AAAGCCCTACAGCAACTGTCG |

**Supplemental Table 3. Gene loading contributions separating cell source and licensing state.**

| Mesenchymal stromal cell |  |  |  |  |  |  |
| --- | --- | --- | --- | --- | --- | --- |
| Gene | PC1 loading coefficient | PC1 vector contribution (%) | PC2 loading coefficient | PC2 vector contribution (%) | PC3 loading coefficient | PC3 vector contribution (%) |
| *HGF* | 0.96466 | 14.55% | -0.07572 | 0.10% | -0.20493 | 1.20% |
| *TNFAIP6* | 0.89026 | 12.39% | 0.10691 | 0.20% | 0.36953 | 3.90% |
| *PDGFA* | -0.86579 | 11.72% | 0.29147 | 1.48% | -0.37888 | 4.10% |
| *CCN2* | -0.85750 | 11.50% | -0.22434 | 0.88% | 0.18885 | 1.02% |
| *ANGPT1* | 0.83951 | 11.02% | -0.25028 | 1.09% | -0.26100 | 1.95% |
| *ACTA2* | 0.64773 | 6.56% | -0.44194 | 3.40% | -0.53426 | 8.16% |
| *THBS1* | -0.61769 | 5.97% | -0.55021 | 5.27% | 0.49708 | 7.06% |
| *CXCL8* | 0.59955 | 5.62% | 0.59082 | 6.08% | 0.42519 | 5.17% |
| *EDN1* | -0.59564 | 5.55% | 0.70595 | 8.68% | 0.30917 | 2.73% |
| *IDO1* | 0.56259 | 4.95% | 0.7689 | 10.30% | 0.27862 | 2.22% |
| *SMAD7* | 0.48626 | 3.70% | 0.71386 | 8.88% | 0.18366 | 0.96% |
| *EDIL3* | -0.45097 | 3.18% | 0.5754 | 5.77% | -0.65598 | 12.30% |
| *SOX9* | -0.28593 | 1.28% | -0.76911 | 10.30% | 0.45579 | 5.94% |
| *TSG101* | -0.22877 | 0.82% | 0.94849 | 15.67% | 0.19493 | 1.09% |
| ANG | -0.20296 | 0.64% | 0.30006 | 1.57% | 0.49708 | 22.43% |
| PDCD1LG2 | -0.13637 | 0.29% | 0.60643 | 6.41% | 0.18885 | 14.79% |
| VEGFA | 0.13306 | 0.28% | -0.89377 | 0.14% | -0.37888 | 4.96% |

**Supplemental Table 4. List of top 20 up- and down-regulated microRNAs in iMSCs**

|  | miRNA | LogFC | Adjusted p-value |
| --- | --- | --- | --- |
| **Upregulated** | **hsa-miR-218-5p** | 2.27 | 7.29E-05 |
|  | **hsa-miR-548n** | 3.12 | 9.61E-03 |
|  | **hsa-let-7f-5p** | 1.64 | 1.03E-02 |
|  | **hsa-miR-7-5p** | 1.90 | 1.41E-02 |
|  | hsa-miR-155-5p | 1.54 | 8.08E-02 |
|  | hsa-miR-543 | 1.34 | 8.44E-02 |
|  | hsa-miR-431-5p | 1.51 | 1.45E-01 |
|  | hsa-miR-370-3p | 1.57 | 2.52E-01 |
|  | hsa-miR-33a-5p | 1.52 | 4.49E-01 |
|  | hsa-miR-30e-5p | 1.51 | 4.49E-01 |
|  | hsa-miR-378f | 1.28 | 4.49E-01 |
|  | hsa-miR-549a | 1.46 | 4.88E-01 |
|  | hsa-miR-329-3p | 1.37 | 4.88E-01 |
|  | hsa-miR-362-5p | 1.44 | 5.16E-01 |
|  | hsa-miR-222-3p | 1.32 | 5.16E-01 |
|  | hsa-miR-299-3p | 1.60 | 5.32E-01 |
|  | hsa-miR-644a | 1.51 | 5.32E-01 |
|  | hsa-miR-148a-3p | 2.12 | 5.37E-01 |
|  | hsa-miR-5196-3p+hsa-miR-6732-3p | 1.56 | 5.80E-01 |
|  | hsa-miR-497-5p | 1.31 | 6.77E-01 |
| **Downregulated** | **hsa-miR-145-5p** | -1.41 | 5.63E-03 |
|  | **hsa-miR-27b-3p** | -1.07 | 1.20E-03 |
|  | hsa-miR-143-3p | -1.31 | 1.77E-01 |
|  | hsa-miR-582-5p | -4.16 | 2.52E-01 |
|  | hsa-miR-181b-5p+hsa-miR-181d-5p | -0.89 | 5.49E-01 |
|  | hsa-miR-137 | -0.70 | 6.98E-01 |
|  | hsa-miR-708-5p | -2.02 | 7.00E-01 |
|  | hsa-miR-450a-5p | -0.56 | 7.00E-01 |
|  | hsa-miR-3161 | -0.98 | 7.01E-01 |
|  | hsa-miR-769-5p | -0.81 | 7.04E-01 |
|  | hsa-miR-363-3p | -0.52 | 7.04E-01 |
|  | hsa-miR-10b-5p | -0.45 | 7.04E-01 |
|  | hsa-miR-4521 | -0.84 | 7.42E-01 |
|  | hsa-miR-542-3p | -0.55 | 7.63E-01 |
|  | hsa-miR-34a-5p | -0.95 | 7.68E-01 |
|  | hsa-miR-593-3p | -0.45 | 7.95E-01 |
|  | hsa-miR-1234-3p | -1.15 | 8.02E-01 |
|  | hsa-miR-424-5p | -0.49 | 8.60E-01 |
|  | hsa-miR-10a-5p | -0.56 | 8.94E-01 |
|  | hsa-miR-196b-5p | -0.43 | 9.32E-01 |

**Supplemental Table 5. Targets of differentially expressed microRNAs in *TGFB1*-iMSCs overlapped with differentially expressed genes tested.** \*Denotes differentially expressed mRNAs between *TGFB1*-iMSCs and MSC(AT). Arrow directions denotes either up- or down-regulated.

| microRNA profiling |  |  |  |  |  |
| --- | --- | --- | --- | --- | --- |
| Gene Target | mRNA in iMSC | miRNA | miRNA in iMSC | Predicted relationship | Observed Relationship |
| ***ACTA2*\*** | ↓ | hsa-let-7a-5p | ↑ | Inhibition | **Observed** |
|  |  | hsa-let-7f-5p | ↑ | Inhibition | **Observed** |
| ***ANGPT1*\*** | ↓ | hsa-miR-145-5p | ↓ | Released inhibition | Not observed |
| ***HGF*\*** | ↓ | hsa-let-7f-5p | ↑ | Inhibition | **Observed** |
|  |  | hsa-miR-27b-3p | ↓ | Released inhibition | Not observed |
| ***EDN1*\*** | ↑ | hsa-let-7f-5p | ↑ | Inhibition | Not observed |
|  |  | hsa-let-7a-5p | ↑ | Inhibition | Not observed |
| ***PDGFA*\*** | ↑ | hsa-let-7f-5p | ↑ | Inhibition | Not observed |
|  |  | hsa-let-7a-5p | ↑ | Inhibition | Not observed |
|  |  | hsa-miR-574-5p | ↑ | Inhibition | Not observed |
| *TNFAIP6* | ↓ | hsa-miR-7-5p | ↑ | Inhibition | **Observed** |
| *CXCL8* | ↓ | hsa-let-7a-5p | ↑ | Inhibition | **Observed** |
|  |  | hsa-let-7f-5p | ↑ | Inhibition | **Observed** |
|  |  | hsa-miR-7-5p | ↑ | Inhibition | **Observed** |
|  |  | hsa-miR-574-5p | ↑ | Inhibition | **Observed** |
| *EDIL3* | ↑ | hsa-miR-27b-3p | ↓ | Released inhibition | **Observed** |
|  |  | hsa-let-7a-5p | ↑ | Inhibition | Not observed |
|  |  | hsa-let-7f-5p | ↑ | Inhibition | Not observed |
|  |  | hsa-miR-7-5p | ↑ | Inhibition | Not observed |
|  |  | hsa-miR-382-5p | ↑ | Inhibition | Not observed |
|  |  | hsa-miR-218-5p | ↑ | Inhibition | Not observed |
| *TSG101* | ↑ | hsa-miR-27b-3p | ↓ | Released inhibition | **Observed** |
|  |  | hsa-let-7a-5p | ↑ | Inhibition | Not observed |
|  |  | hsa-let-7f-5p | ↑ | Inhibition | Not observed |
|  |  | hsa-miR-7-5p | ↑ | Inhibition | Not observed |

**Supplemental Table 6. Loading contributions of MSC-induced HUVEC tube formation.**

| **HUVEC Tube Formation** |  |  |  |  |  |  |
| --- | --- | --- | --- | --- | --- | --- |
| **Parameters** | **PC1 loading coefficient** | **PC1 vector contribution (%)** | **PC2 loading coefficient** | **PC2 vector contribution (%)** | **PC3 loading coefficient** | **PC3 vector contribution (%)** |
| **Total branches length** | 0.98877 | 8.29 | -0.02841 | 0.02 | -0.00043 | 0.00 |
| **Nb meshes** | 0.98765 | 8.28 | 0.0096 | 0.00 | -0.01740 | 0.02 |
| **Nb Master Junction** | 0.97657 | 8.09 | 0.0573 | 0.09 | 0.12535 | 0.90 |
| **Nb junctions** | 0.96753 | 7.94 | 0.05338 | 0.08 | 0.12706 | 0.92 |
| **Nb branches** | 0.96561 | 7.91 | -0.11700 | 0.39 | 0.16768 | 1.61 |
| **Total master segment lengths** | 0.94955 | 7.65 | -0.20897 | 1.26 | 0.21266 | 2.59 |
| **Nb Master segments** | 0.93331 | 7.39 | -0.10837 | 0.34 | 0.15241 | 1.33 |
| **Nb Segments** | 0.92929 | 7.33 | 0.24555 | 1.74 | 0.23195 | 3.08 |
| **Total segments length** | 0.91538 | 7.11 | 0.35871 | 3.71 | -0.13106 | 0.98 |
| **Total branching length** | 0.86547 | 6.35 | 0.46227 | 6.16 | 0.00663 | 0.00 |
| **Total Mesh area** | 0.84176 | 6.01 | -0.44009 | 5.58 | 0.28818 | 4.75 |
| **Nb Pieces** | 0.75904 | 4.89 | -0.24625 | 1.75 | -0.22593 | 2.92 |
| **Mesh index** | 0.69214 | 4.06 | -0.61009 | 10.73 | 0.18942 | 2.05 |
| **Total length** | -0.62529 | 3.32 | 0.07422 | 0.16 | 0.73649 | 31.03 |
| **Branching interval** | -0.51704 | 2.27 | 0.34696 | 3.47 | 0.65675 | 24.68 |
| **Nb isolated segments** | 0.49098 | 2.05 | 0.82987 | 19.86 | -0.13638 | 1.06 |
| **Total isolated branches length** | 0.32179 | 0.88 | 0.8533 | 20.99 | -0.29286 | 4.91 |
| **Nb Nodes** | -0.14143 | 0.17 | 0.84587 | 20.63 | 0.48036 | 13.20 |
| **Mean mesh size** | 0.03074 | 0.01 | 0.32368 | 3.02 | -0.26340 | 3.97 |

**Supplemental Table 7. Gene loading contributions from MSC-treated monocyte/macrophages.**

| **Monocyte/macrophage polarization** |  |  |  |  |  |  |
| --- | --- | --- | --- | --- | --- | --- |
| **Gene** | **PC1 loading coefficient** | **PC1 vector contribution (%)** | **PC2 loading coefficient** | **PC2 vector contribution (%)** | **PC3 loading coefficient** | **PC3 vector contribution (%)** |
| ***CCR7*** | 0.96483 | 17.82 | 0.02601 | 0.02 | -0.06051 | 0.36 |
| ***IL12A*** | 0.91614 | 16.07 | 0.12068 | 0.41 | 0.12302 | 1.48 |
| ***IL1B*** | 0.64152 | 14.75 | 0.06126 | 0.11 | 0.33266 | 10.83 |
| ***TREM1*** | 0.79091 | 11.98 | 0.33638 | 3.22 | 0.20962 | 4.30 |
| ***IL10*** | 0.78899 | 11.92 | 0.4267 | 5.18 | -0.16532 | 2.67 |
| ***HLA-DRA*** | 0.64152 | 7.88 | 0.43963 | 5.50 | -0.16682 | 2.72 |
| ***PDL1*** | 0.63313 | 7.67 | 0.68773 | 13.47 | 0.12439 | 1.51 |
| ***HMOX1*** | 0.54265 | 5.64 | 0.63065 | 11.32 | -0.38063 | 14.17 |
| ***CD206*** | 0.48770 | 4.55 | 0.2996 | 2.56 | 0.65802 | 42.36 |
| ***STAB1*** | 0.28908 | 1.60 | 0.84652 | 20.40 | -0.27932 | 7.63 |
| ***CD163*** | 0.07683 | 0.11 | 0.87551 | 21.82 | 0.05104 | 0.25 |
| ***CD86*** | 0.02262 | 0.01 | 0.74932 | 15.98 | 0.34604 | 11.71 |

**Supplemental Table 8. Euclidean distances to resolve inter-condition differences.**

| Experiment | Samples | $\Delta PC1$ | $(\Delta PC1)^2$ | $\Delta PC2$ | $(\Delta PC2)^2$ | Euclidean distance (ED) |
| --- | --- | --- | --- | --- | --- | --- |
| **Nanostring Gene Expression (Fig. 2c)** | iMSC LIC – UNL | 0.763 | 0.583 | 3.530 | 12.460 | **3.612** |
|  | AT1 LIC – UNL | 3.178 | 10.101 | 3.845 | 14.784 | **4.988** |
|  | AT4 LIC – UNL | 2.919 | 8.518 | 3.336 | 11.131 | **4.433** |
| **microRNA Profiling (Fig. 3a)** | iMSC – AT1 | -0.611 | 0.373 | 1.784 | 3.183 | **1.886** |
|  | iMSC – AT2 | -21.247 | 451.415 | -5.758 | 33.154 | **22.013** |
|  | iMSC – AT3 | 10.147 | 102.953 | -9.933 | 98.658 | **14.199** |
|  | iMSC – AT4 | 0.704 | 0.495 | 3.494 | 12.205 | **3.564** |
| **Angiogenic Tube Formation (Fig. 4c)** | iMSC – POS | -3.373 | 11.376 | -2.051 | 4.205 | **3.947** |
|  | iMSC – NEG | 4.076 | 16.616 | 0.243 | 0.059 | **4.084** |
|  | iMSC – AT1 | 2.445 | 5.977 | -3.181 | 10.122 | **4.012** |
|  | iMSC – AT2 | -2.128 | 4.529 | -3.804 | 14.471 | **4.359** |
|  | iMSC – AT3 | 3.153 | 9.944 | -3.259 | 10.621 | **4.535** |
|  | iMSC – AT4 | -2.487 | 6.184 | -3.010 | 9.062 | **3.905** |
| **Immunomodulatory Monocyte Co-culture (Fig. 5d)** | iMSC-Mono | -4.371 | 19.103 | 3.077 | 9.467 | **5.345** |
|  | AT1-Mono | -4.449 | 19.796 | 0.503 | 0.253 | **4.478** |
|  | AT2-Mono | -3.806 | 14.487 | 1.946 | 3.788 | **4.275** |
|  | AT3-Mono | -2.322 | 5.391 | 0.533 | 0.284 | **2.382** |
|  | AT4-Mono | 0.443 | 0.197 | 1.915 | 3.668 | **1.966** |
|  | iMSC-AT1 | 0.079 | 0.006 | 2.574 | 6.626 | **2.575** |
|  | iMSC-AT2 | -0.564 | 0.319 | 1.131 | 1.278 | **1.264** |
|  | iMSC-AT3 | -2.049 | 4.198 | 2.544 | 6.473 | **3.267** |
|  | iMSC-AT4 | -4.814 | 23.176 | 1.162 | 1.349 | **4.952** |
