## Supplemental Figures for "*TGFB1*-Engineered Induced Mesenchymal Stromal Cells Exhibit High Immunomodulatory Potency and Durably Reprogram Synovial Macrophages in Osteoarthritis"

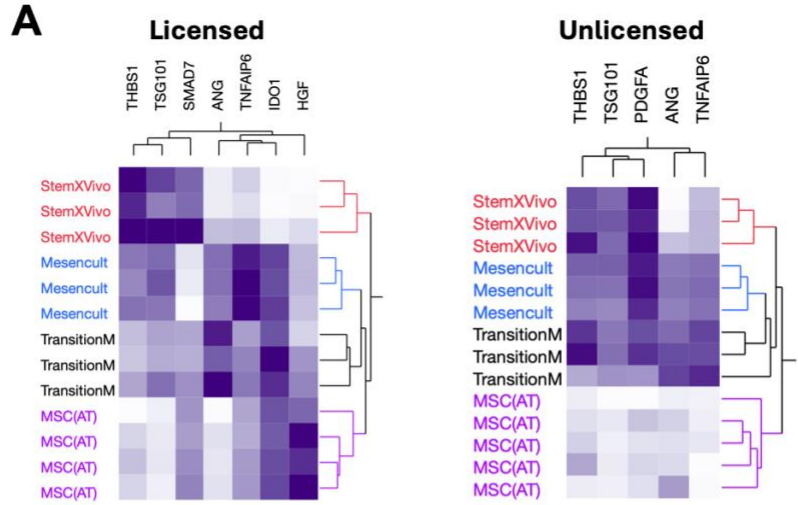

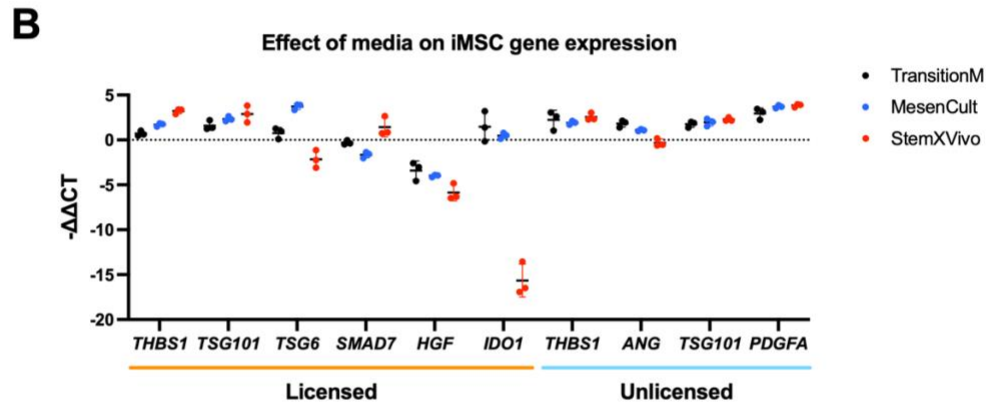

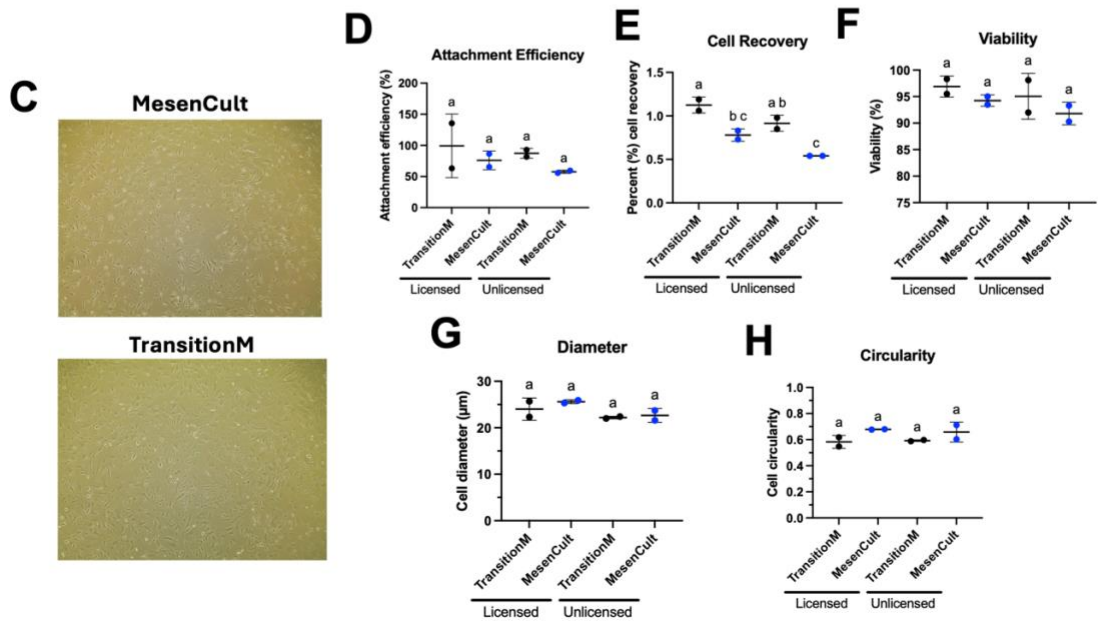

**Supplementary Figure 1. Selection of iMSC culture medium based on gene expression and cell characteristics.** iMSCs were expanded in StemXVivo, MesenCult™-ACF Plus (MesenCult), or a transition protocol in which cells were initiated in StemXVivo and switched to MesenCult™-ACF Plus (TransitionM) and compared against primary MSC(AT) expanded in MesenCult™-ACF Plus. (A) Unsupervised hierarchical clustering of marker gene expression in unlicensed and licensed (pro-inflammatory cytokine-stimulated) cells. (B) Marker gene expression ($-\Delta\Delta\text{Ct}$) across media for licensed (THBS1, TSG101, TSG6, SMAD7, HGF, IDO1) and unlicensed (THBS1, ANG, TSG101, PDGFA) panels; dotted line indicates the MSC(AT) reference. (C) Representative phase-contrast images of iMSCs in MesenCult and TransitionM. (E) Attachment efficiency at 4 h, (F) cell recovery at 24 h, (G) viability, (H) cell diameter, and (I) cell circularity in TransitionM and MesenCult, under licensed and unlicensed conditions. Data are mean ± SD of $n = 2$ or 3 independent biological replicates; individual replicates are shown. Groups not sharing a letter differ significantly.

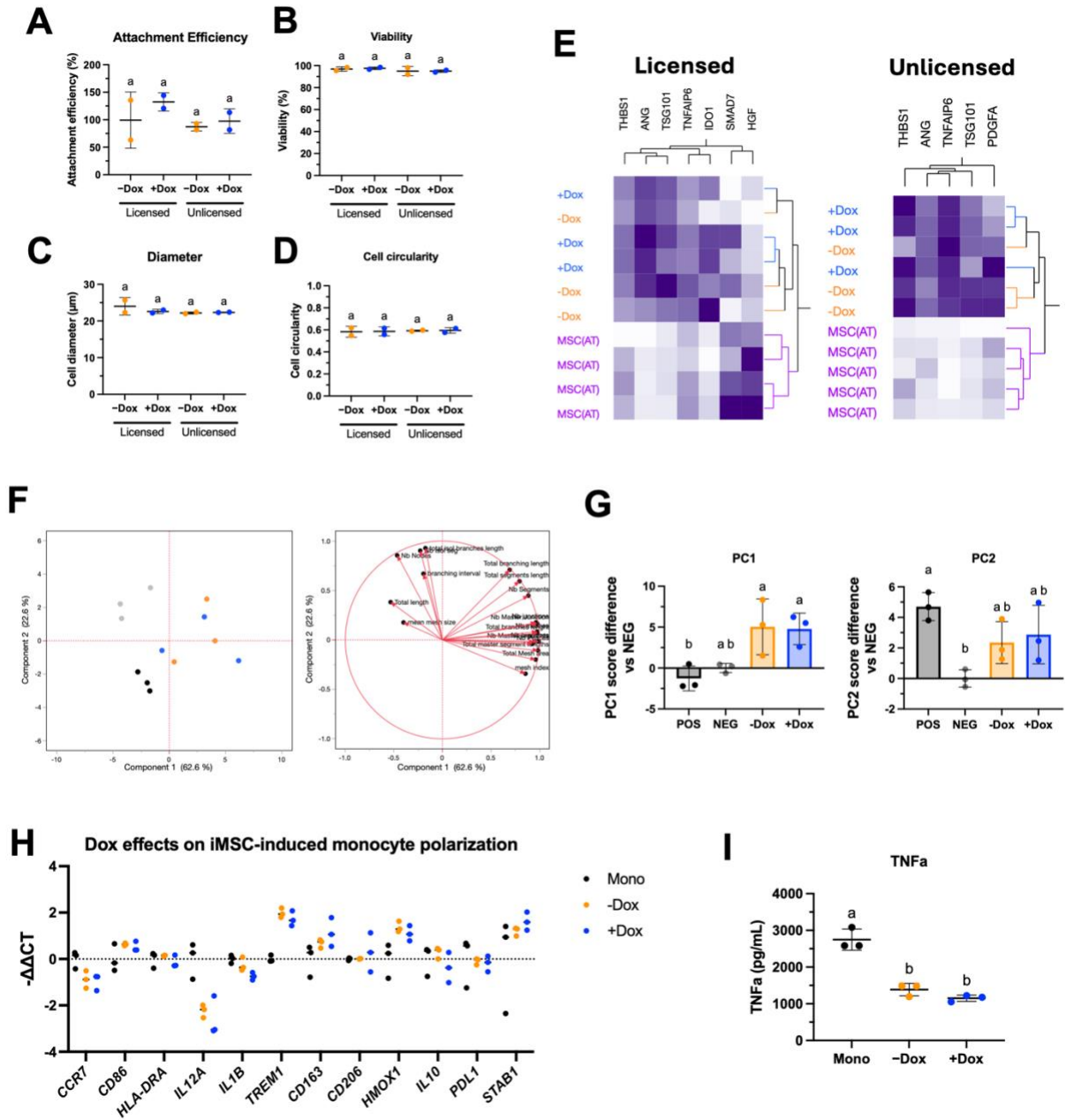

**Supplementary Figure 2. Doxycycline induction does not alter *TGFβ1*-iMSC culture characteristics, marker expression, or immunomodulatory function.** *TGFβ1*-iMSCs were cultured in TransitionM and assessed in the absence (-Dox) or presence (+Dox) of doxycycline. (A) Unsupervised hierarchical clustering of licensed (left) and unlicensed (right) gene expression in ±Dox iMSCs and primary MSC(AT). (B) Attachment efficiency at 4 h, (C) viability, (D) cell diameter, and (E) cell circularity in licensed and unlicensed *TGFβ1*-iMSCs ±Dox. (F) PCA of tube formation readouts from ±Dox *TGFβ1*-iMSC-treated HUVECs. Score plot (left) and loading plot (right). (G) PC1 and PC2 score differences relative to the negative control. (H) Monocyte polarization marker expression ($-\Delta\Delta Ct$ relative to the monocyte-alone control) following co-culture with ±Dox *TGFβ1*-iMSCs. (I) TNFα secretion (pg/mL) by monocytes cultured alone (Mono) or co-cultured with ±Dox *TGFβ1*-iMSCs. Data are the mean ± SD of $n = 3$ independent biological replicates; individual replicates shown.

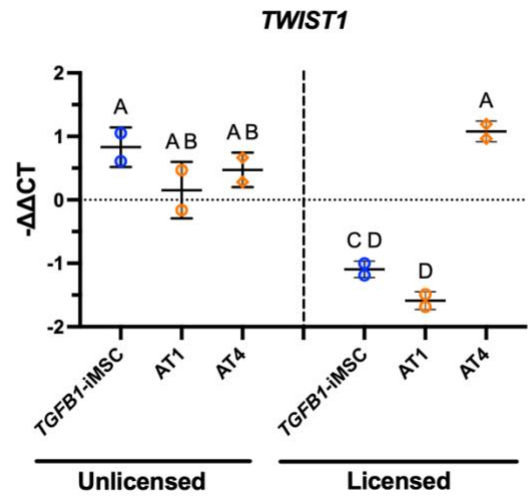

**Supplemental Figure 3. *TWIST1* expression in iMSCs and MSC(AT) under unlicensed and licensed conditions.** RT-qPCR measured *TWIST1* expression in TGFB1-iMSCs (blue) and adipose tissue-derived MSCs from two donors, AT1 and AT4 (orange), under unlicensed (left) and licensed (right) conditions. The dotted line marks no change relative to the reference. Data are mean ± SD ($n = 2$ biological replicates). Groups with different letters differ significantly ($p < 0.05$, two-way ANOVA with Tukey's post hoc test).

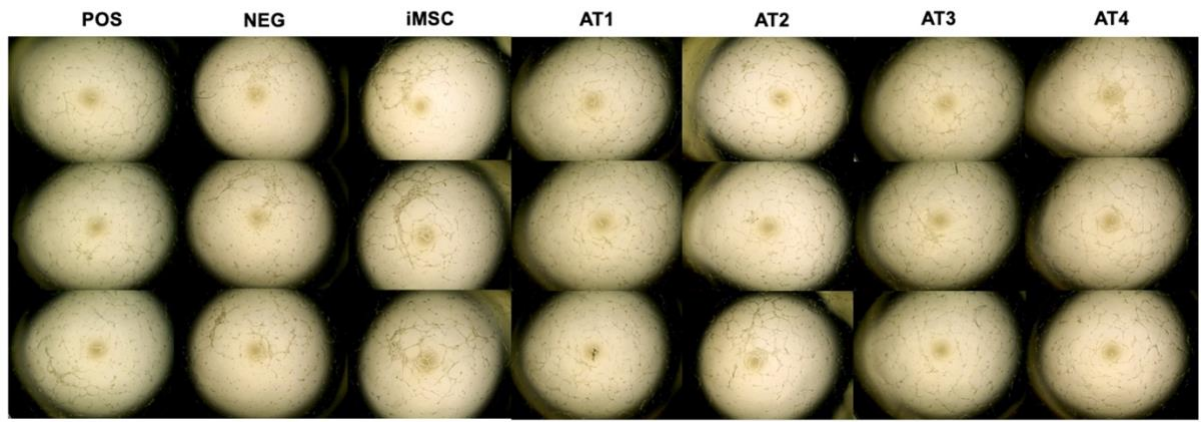

**Supplemental Figure 4: HUVEC tube formation images.** Representative brightfield images of HUVEC tube formation 6 hours after treatment with positive control (POS), negative control (NEG), iMSC-conditioned media, or MSC(AT) donor (AT1–AT4)-conditioned media. $n = 3$ independent wells per condition.

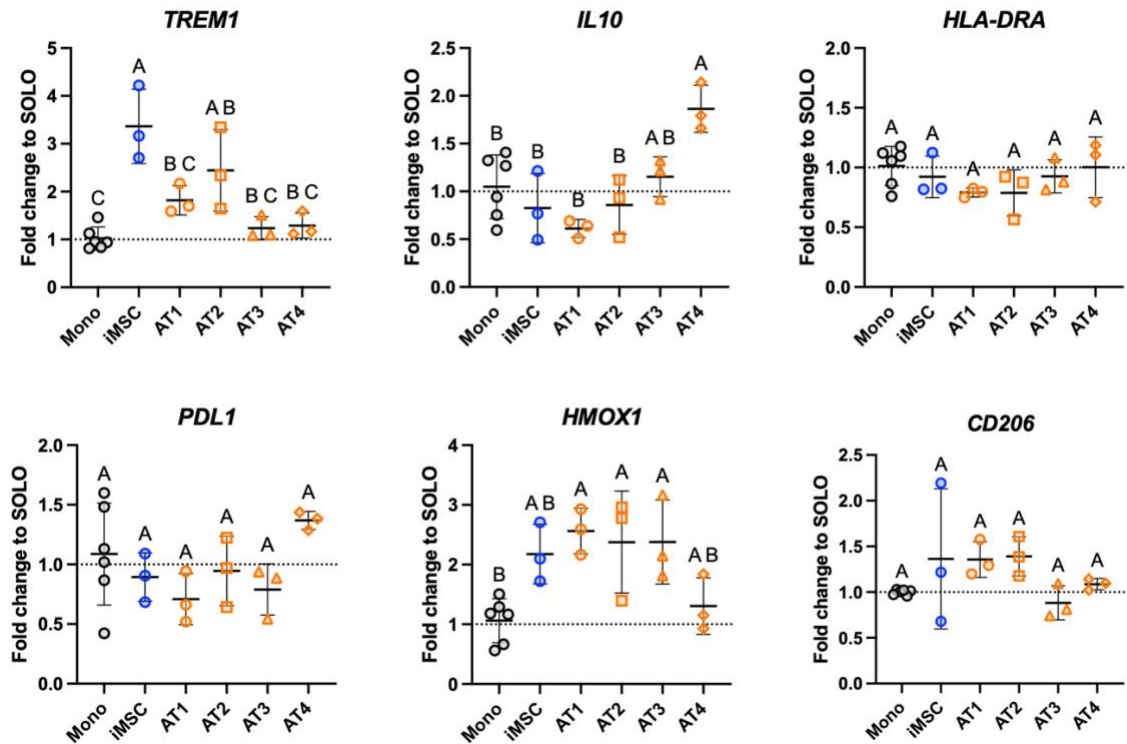

**Supplemental Figure 5: Monocyte/macrophage polarization gene expression following co-culture with iMSCs and MSC(AT) donors.** (A) Fold change relative to SOLO across a broader panel of pro- and anti-inflammatory polarization genes (CCR7, CD86, HLA-DRA, IL12A, IL1B, TREM1, CD163, CD206, HMOX1, IL10, PDL1, STAB1) in macrophages co-cultured with iMSCs or MSC(AT) donors (AT1–AT4). (B) Individual gene expression fold changes relative to SOLO for TREM1, IL10, HLA-DRA, PDL1, HMOX1, and CD206. Data are presented as mean ± SD, $n = 3$ independent experiments. We performed statistical comparisons using one-way ANOVA with Tukey's post hoc test; groups sharing a letter are not significantly different ($p > 0.05$).
